# Hydrodynamic dispersion drives viral-cellular contact for gene delivery in porous media

**DOI:** 10.64898/2026.01.15.699726

**Authors:** Vishal Srikanth, Micah Mallory, Andrew J. Ulmer, Wesley R. Legant, Andrey V. Kuznetsov, Yevgeny Brudno

## Abstract

Reactive biological processes often hinge on rare collisions between particles that occupy vastly different physical regimes, yet the transport physics that govern these encounters remain poorly understood. Biological cell-virus encounters offer a uniquely quantifiable instance of this general problem: collisions between particles whose transport is governed by entirely different physical mechanisms, yet whose interactions determine system-level function. In stagnant liquids, nanoscale viral vectors explore space only through slow Brownian diffusion, while microscale cells rapidly sediment, producing species separation that suppresses the virus-cell interaction interface. Here we show that liquid absorption into a dry, macroporous sponge enhances viral-cellular interactions by shifting the system into an advection-dispersion regime that circumvents this sedimentation-diffusion limit. By integrating experimental results with a multiscale simulation model, we demonstrate that the tortuous sponge porosity converts capillary-driven flow into convective mixing, driving orders-of-magnitude increases in viral-cellular collision rates. Coupling these dispersive transport dynamics with a probabilistic capture model reveals that hydrodynamic dispersion accounts for the multifold enhancement in viral-cellular transduction efficiency observed in porous sponges. These results provide a quantitative framework for emergent collision dynamics in complex porous media and establish a generalizable strategy to optimize active transport in spatiotemporally heterogeneous biological systems.

## 1 Introduction

Many biological and chemical processes hinge on the encounter between particles that occupy vastly different physical regimes, yet the transport physics that governs these interactions remain incompletely understood. Viral transduction – where nanoscale vectors must locate and bind to microscale cells – provides a particularly illustrative and experimentally tractable example of this broader problem. In therapeutic settings, this process underlies the ability to engineer cellular function by introducing new genetic material, enabling transformative modalities such as CAR T-cell therapies^1–4^, which are a potent gene therapy for treating various types of cancer^5^. Despite their clinical promise, the physical step of viral transduction – delivery of genetic cargo to the cell – remains a critical manufacturing bottleneck that severely limits patient access to these life-saving treatments^6–9^. Improving the efficiency of this step holds the key to making these transformative therapies more scalable, affordable, and widely available.

The inefficiency of passive cell transduction stems from a fundamental physical challenge. In a typical liquid suspension, microscale cells and nanoscale viruses are initially homogeneously distributed throughout the liquid volume. The kinetics of these particles are governed by different mechanisms of disparate scales, leading to their physical separation and severely limiting the number of cell-virus collisions that occur in the static medium. Furthermore, since a virus must locate and bind to a specific, sparsely distributed receptor on the cell surface, a single cell-virus contact is often insufficient for gene transfer, and the virus needs to probe the cell membrane across multiple collisions to achieve successful binding^10–13^.

Various methods have been developed to force cell-viral interaction and catalyze transduction, either by mobilizing the cells and viruses using microfluidic devices^14–18^ or low-speed centrifugation (spinoculation)^19,20^, by improving viral-cellular binding through chemical reagents such as polybrene^21^ or retronectin^22^, or by transiently increasing membrane permeability using mechanoporation^23^. However, these approaches rely on specialized equipment, expensive reagents, or suffer from limited throughput, making them impractical for large-scale manufacturing. Moreover, because the physical mechanisms underlying their enhanced efficiency are poorly defined, active cell transduction approaches remain difficult to predict or generalize, limiting their translation into robust clinical workflows. Thus, despite these advances and in spite of its fundamental flaws, transduction in stagnant liquid remains the clinical standard for CAR T cell manufacturing.

In previous work, we reported a fundamentally different approach to enhancing viral transduction that bypasses the need for specialized devices or reagents, relying instead on the intrinsic physical properties of biomaterials. We reported that dry, macroporous biomaterial sponges can achieve highly efficient transduction with an intriguingly simple workflow^24,25^. When a liquid droplet holding a suspension of cells and viruses was kept stagnant, we observed poor transduction efficiency (°10%). However, simply adding the same liquid droplet to the dry sponge and allowing it to absorb into the pores increases transduction efficiency to >80%, a multifold improvement. Notably, this enhancement was observed across a wide range of macroporous biomaterials^26^ and extended to the delivery of lipid nanoparticles, suggesting that the effect arises from general transport and interaction physics rather than material- or biology-specific factors. The stark contrast between the simplicity of the process and the magnitude of the improvement raises a fundamental question: what physical mechanisms underpin this remarkable enhancement?

Initial studies pointed to liquid absorption and macroporosity as key factors behind sponge-enhanced transduction^24^. Transduction was not enhanced either in pre-wetted sponges, which decrease the absorption rate, or in sponges with nano-sized pores, which arrest cell advection. Further investigation revealed that sponge-enhanced transduction efficiency was closely correlated to the rate of fluid flux during absorption^27^. These observations strongly suggested that sponge transduction enhancement is driven by the physical process of liquid flow through the confined sponge porosity. Indeed, these physical mechanisms are known to improve particle interaction rates in other systems, including in microfluidic devices, which leverage advection and colocalization to enhance transduction^15–17^ and the use of shear flow to increase particle collision frequency in bacterial conjugation^28^. In principle, a porous sponge might combine these principles: rapid, hygroscopic absorption generates strong convective advection, the interaction of the absorbing liquid with the pore walls creates hydrodynamic shear, and the pore architecture itself serves to colocalize particles. This led us to hypothesize that the sponges fundamentally alter the physics of cell-virus collisions, transforming the system from a passive, diffusion-limited regime to an active, convection-dominated one.

We now report a comprehensive physical model that is tightly integrated with experimental measurements to deconstruct the mechanisms of sponge-mediated transduction. By simulating cell-virus collisions under therapeutically relevant conditions, we directly contrast the cell-virus kinetics within a porous sponge against those occurring within a stagnant droplet. This comparative analysis allows us to isolate and quantify the contributions of the dominant physical forces driving transduction enhancement, from convective particle dispersion during absorption to particle colocalization within the pores. By establishing how stochastic porous media reshape multiscale particle transport and reaction kinetics, this work reveals the physical principles that govern collision-limited processes in these complex environments. In doing so, it provides not only a predictive framework for optimizing biomaterial design and manufacturing workflows, but also a model system for understanding emergent transport phenomena that arise whenever flow, geometry, and stochasticity interact across scales.

## 2 Results

### 2.1 Analysis of viral cellular transduction in a stagnant liquid droplet

Viral genetic modification in a stagnant liquid medium remains the dominant paradigm for clinical and preclinical cell manufacturing^29^, including current lentiviral transduction protocols used for CAR T-cell production. In these settings, cells and viral vectors are typically co-incubated in static wells, bags, or bioreactors, where productive gene transfer depends entirely on the frequency and outcome of stochastic cell–virus encounters. Understanding the physical limits of this stagnant-liquid regime is therefore a necessary prerequisite for evaluating and rationally improving any alternative transduction strategy. To quantitatively understand how cell-virus collisions translate into transduction events under clinically relevant conditions, we developed a simulation model of transduction in a stagnant liquid droplet that closely mirrors standard experimental protocols. We simulated a representative sub-volume of the liquid droplet, tracking individual cell and virus trajectories using a discrete particle model (**figure 1a**). The domain height was set by the equilibrium droplet height for a given liquid volume estimated analytically from the droplet shape^30^ (**supp. figure 1**), while the lateral width was fixed at 1 mm. To create a physically representative, computationally tractable system, we imposed periodic boundary conditions in the lateral directions, a reasonable approximation because the cell motion is dominated by vertical gravitation sedimentation, while the root-mean-square displacement of viruses over the simulation time (°0.14 mm) remains small relative to the domain size. Reflecting wall boundary conditions were applied at the liquid-air and liquid-solid interfaces, modeling the inability of cells or viruses to cross the droplet boundary due to surface tension^31^. Particle trajectories and cell-virus collisions were tracked for 45 minutes, corresponding to the full experimental transduction window.

**Figure 1.**
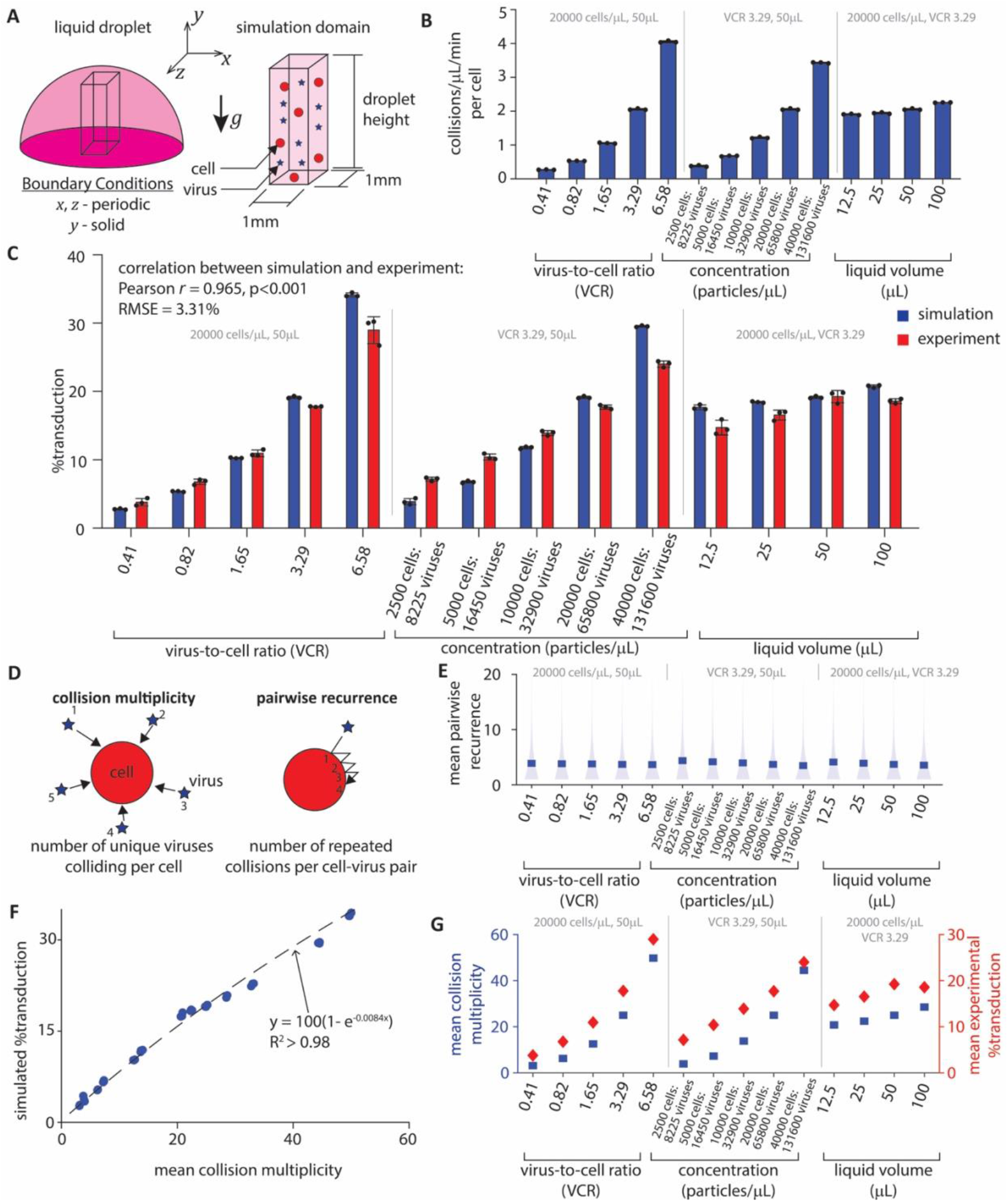
(a) Simulation geometry and boundary conditions used to model transduction in a stagnant liquid droplet. (b) The simulation model predicts cell-virus collisions occurring over time for various liquid volumes, virus-to-cell ratios, and cell-virus concentrations. The constant parameters in each single parameter variation are a concentration of 20,000 cells/µL, VCR of 3.29, and liquid volume of 50µL. (c) A probabilistic transduction model predicts viral-cellular transduction outcome from cell-virus collisions. The simulation model predicts transduction efficiency for various VCRs, cell-virus concentrations, and liquid volumes. (d) Cell-virus collisions are further analyzed by calculating collision multiplicity and pairwise recurrence. (e) The model shows that pairwise recurrence is unchanged across all parameter variations, (f-g) whereas the improvement in transduction efficiency is brought by the increase in collision multiplicity. The blue markers show simulated mean pairwise recurrence and collision multiplicity, and the shaded blue areas show the probability distribution of simulated pairwise recurrence. The red marker shows experimental transduction efficiency.

We evaluated our model across experimentally relevant conditions spanning variations in virus-to-cell ratio, absolute cell-virus concentration, and total droplet volume. The model revealed that the cell-virus collision frequency – defined as the number of collisions per unit volume per unit time – increases strongly with both the virus-to-cell ratio (VCR) and the cell-virus concentration but increases only marginally with liquid volume (**figure 1b**). This parametric dependence confirmed that collision kinetics in stagnant liquid droplets are controlled primarily by the local availability of cell-virus pairs. Consistent with this interpretation, the simulated cell-virus collision frequency per cell was strongly correlated with the experimentally measured transduction efficiency, establishing collision rate as the dominant determinant of transduction outcomes in stagnant liquid droplets. This behavior is expected in a closed system over the 45-minute transduction window, where cell proliferation is negligible^32^, and viral transport is diffusion limited. Under these conditions, collision rate serves as an effective proxy for the extent of viral exploration of the cell surface, integrating membrane sampling, receptor engagement, and the probability of successful cell reprogramming^13^.

This link allowed us to introduce a minimal probabilistic model relating cell-virus collisions to transduction. The model is defined by a single transduction probability, *p*_*TE*_, that maps individual cell-virus collisions to the likelihood of successful cell reprogramming events. Viral transduction is initiated by receptor-mediated binding between the viral envelope and cognate receptors on the cell, triggering internalization and cytoplasmic release of the genetic cargo. Subsequent reverse transcription, nuclear transport, and genomic integration complete the reprogramming process. The parameter *p*_*TE*_ therefore captures the combined probability of these downstream biological processes – receptor engagement, internalization, and successful genome integration – to predict transduction following collision. Using Bernoulli trials applied to the simulated cell-virus collision history, we numerically estimated *p*_*TE*_ = 0.0023 (95% CI [0.0010, 0.0043]) via Markov Chain Monte Carlo (MCMC), calibrated solely from the experiments varying the cell-virus concentration. This single calibrated parameter was then held fixed to predict transduction efficiency across all remaining experimental conditions (**figure 1c; supp. table 1**). The model accurately captured that observed increases in transduction efficiency with VCR and with absolute cell-virus concentrations, while predicting only weak dependence on liquid volume – consistent with collision-limited transport in a static medium. At liquid volumes exceeding 50 µL, the simulation predicted a continued increase in transduction efficiency, whereas experiments showed a decline. Our experiments reveal a decline in cell viability at these higher liquid volumes, likely due to excessive cell crowding, impaired proliferation and nutrient depletion once cells sediment at the bottom surface (**supp. figure 2**)^33^. Since poor viability directly compromises cellular function, this effect plausibly accounts for the deviation between model and experiments at large liquid volumes for these high cell concentrations.

#### Transduction in a stagnant liquid droplet is inefficient due to gravitational separation

Despite measurable cell-virus collisions, transduction in a stagnant liquid droplet remains inefficient, suggesting a mismatch between collision kinetics and productive biological outcomes. To understand the physical drivers behind these trends, we analyzed the nature of the cell-virus collisions using two complementary metrics (**figure 1d**): collision multiplicity, defined as the number of distinct viruses that collide with an individual cell, and pairwise recurrence, defined as the number of times a given cell-virus pair collides. We found that pairwise recurrence remained constant across all conditions (**figure 1e**), consistent with its dependence on Brownian diffusivity, which is fixed by media viscosity, temperature, and particle size. As a result, pairwise recurrence cannot be readily modified without altering fluid properties, incubation conditions, particle dimensions, or by imposing strong geometric confinement using microfluidic chambers, which can be detrimental to viral stability or cell proliferation^29,34–36^.

In contrast, change in transduction efficiency tracked strongly with collision multiplicity (**figure 1f&g**), identifying it as the dominant physical driver of productive gene transfer. Our model predicts that transduction is driven by the cumulative probability of binding across multiple encounters. In stagnant, unconfined droplets, viral particles are not restricted to the proximity of any individual cell. Thus, once diffusive escape occurs, further collisions between the same cell-virus pair become unlikely. Consequently, successful transduction in unconfined media typically requires the cell to encounter multiple distinct viral particles to reach the necessary binding threshold. In a stagnant liquid droplet, collision multiplicity can therefore be increased only by supplying more viral particles (higher VCR) or by reducing particle separation (higher particle concentration), approaches that raise efficiency empirically but leave the underlying transport limitation unresolved.

To understand how transport dynamics evolve during transduction in a stagnant liquid, we examined the spatiotemporal distribution of cell-virus collisions predicted by our model. The simulation revealed two distinct dynamical phases: a transient “settling phase,” followed by a prolonged “static phase” (**figure 2a**). Initially, cells are uniformly distributed, but due to their negative buoyancy they settle to the bottom of the droplet over an average time period of °15 minutes. In contrast, nanoscale viruses have terminal velocities over three orders of magnitude smaller and are subject to significant Brownian motion, allowing them to remain suspended throughout the liquid for the entire transduction period. Analysis of collision histories revealed that a majority of both collisions and successful transduction events (50-75%) occur during the transient settling phase, which is characterized by elevated collision multiplicity (**figure 2b&c**). As cells accumulate at the bottom of the droplet, local cell concentration increases and the limited viruses in that vicinity are quickly depleted. Consequently, the local VCR at the bottom of the droplet decreases by over two orders of magnitude over the course of the experiment (**figure 2d**). Meanwhile, the vast majority of viruses remain suspended in the upper portion of the droplet, unable to diffuse to the settled cells (**supp. figure 3b**). Slow viral diffusion combined with the low viral particle stability at physiological temperatures^37^ means most viral particles will degrade before they encounter a cell. To quantify virus arrival at the settled cells over extended periods of the static phase, we theoretically simulated the diffusion of viral particles toward the settled cells by assuming various values of the viral particle half-life (1-8h) that modeled clinically relevant viral degradation typically observed for retro- and lentiviral vectors^37–40^. We concluded that less than 15% of viral particles reach the settled cells in a 50µL droplet even over a 10-hour duration of the static phase due to the large particle separation distance (**supp. figure 3c**). These results identify gravitational settling, rather than Brownian diffusion, as the dominant physical mechanism limiting transduction in a stagnant liquid droplet. This conclusion was further supported by a simulation in which gravitational settling was disabled, yielding a sustained increase in transduction efficiency over time and directly demonstrating the impact of gravity-driven particle separation (**supp. figure 4**).

**Figure 2.**
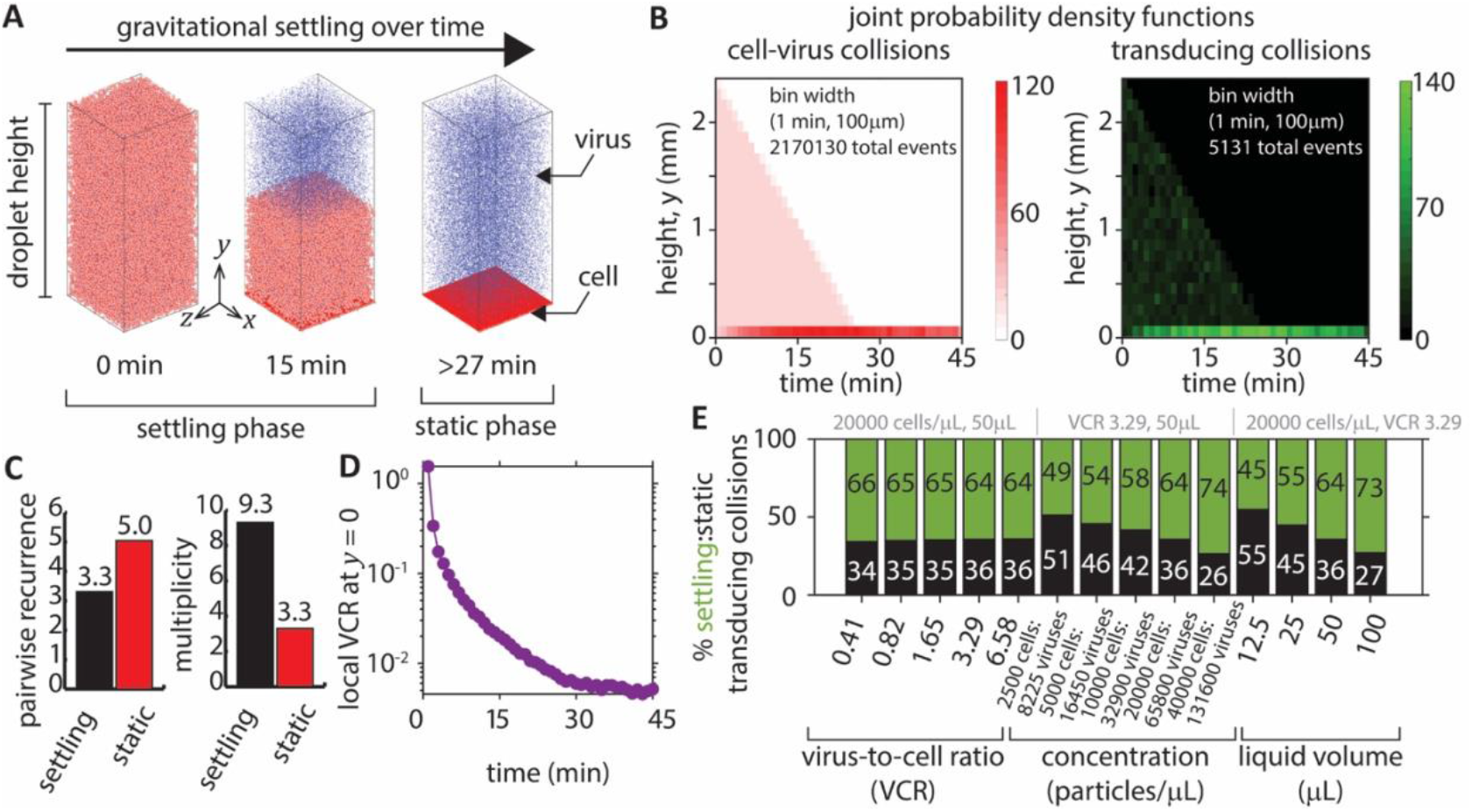
(a) Distributions of the cell and viral concentrations plotted over time show that the settling of cells separates the cell and viral particles – dividing the process into settling and static phases. (b) Probability densities of both cell-virus collisions and successful transduction show the settling process followed by transduction localization at the bottom of the droplet (at *y* = 0), i.e. the static phase. (c) Marginally higher pairwise recurrence in the settling phase is outweighed by a dramatic decrease in collision multiplicity. Lower local VCR is observed during the static phase compared to the settling phase due to cell settling and viral binding over time. Plots (a)-(c) are made for a cell-virus concentration of 20000 cells/µL, liquid volume of 50µL, and VCR = 1.65. (e) Consequently, the model predicts that a majority of transducing collisions (50-80%) occur during the settling phase, whereas a small number of transductions occur during the static phase. Percentages of transducing collisions occurring in the settling phase are shown in green and static phase in black. The constant parameters in each single parameter variation are a concentration of 20,000 cells/µL, VCR of 3.29, and liquid volume of 50µL.

### 2.2 Developing a model of liquid absorption and cell-virus kinetics in porous sponges

#### Convection and colocalization mechanisms are the primary hypotheses for sponge transduction enhancement

The analysis above establishes that stagnant-liquid transduction is fundamentally limited by gravity-driven particle sedimentation and the resulting collapse of collision multiplicity. Thus, strategies to improve transduction must actively reshape transport to sustain encounters between cells and new viral particles over time. Porous biomaterial sponges introduce such a regime. When a suspension of cells and viruses is absorbed by a dry macroporous sponge, transduction efficiency increases dramatically relative to a stagnant droplet (**figure 3a**) revealing an unexpectedly strong coupling between material architecture and transport dynamics^27^. Sponge-mediated transduction enhancement was strongly correlated with the flux of liquid absorption, with transduction increasing from 24% to over 50% under identical cell-virus concentration and sponge compositions. Transduction efficiency was also correlated with pore size, with smaller pores increasing efficiency from 50% to over 80%. These observations suggest liquid absorption into a stochastic pore network as a previously underappreciated driver of cell-virus kinetics and, ultimately, cell transduction. We hypothesize that this enhancement arises from two physical mechanisms intrinsic to porous absorption: (1) convective dispersion of cells and viruses as liquid is absorbed through the sponge's stochastic pore network, and (2) sustained cell/virus colocalization within the sponge's pores following absorption.

**Figure 3.**
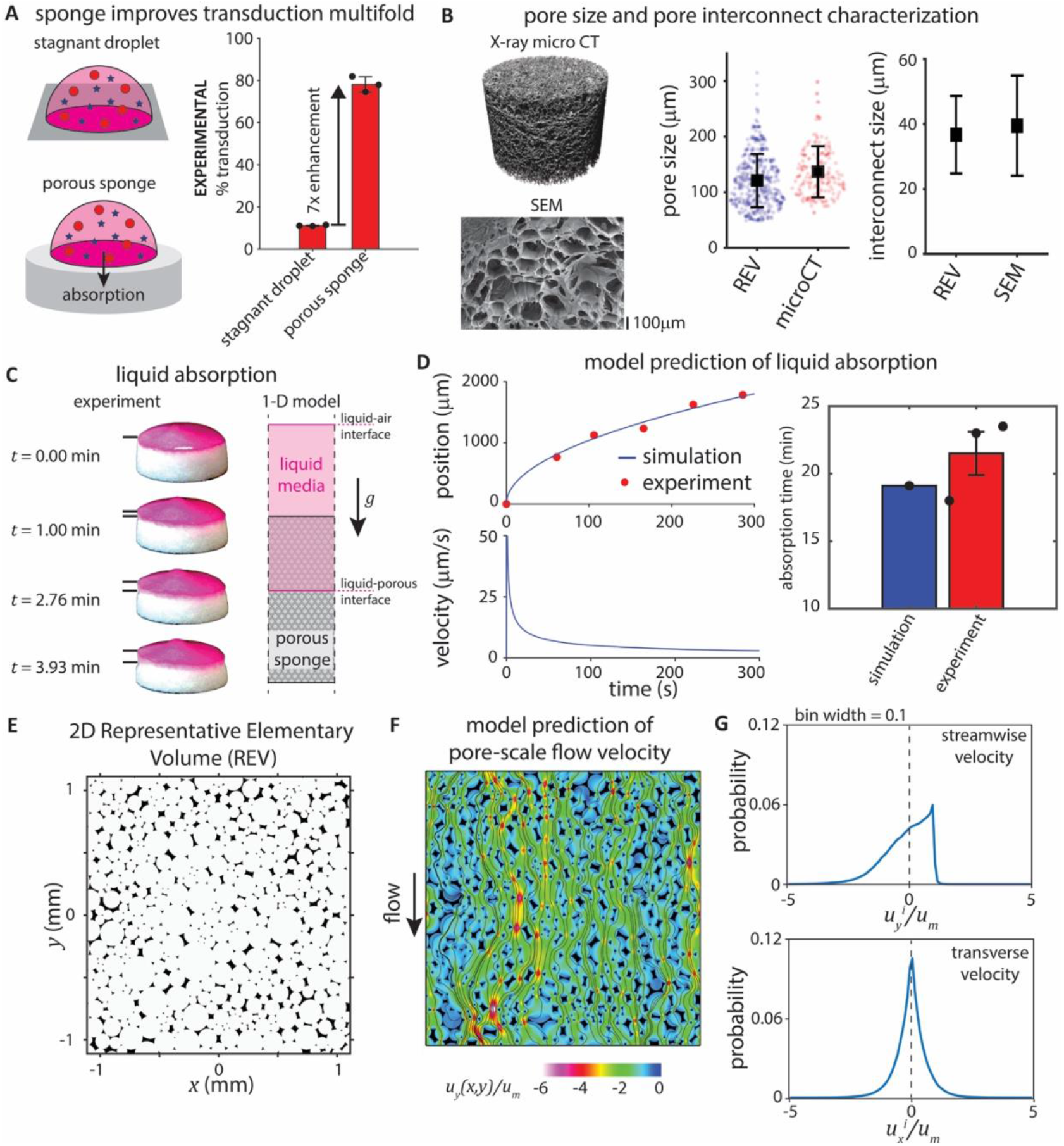
(a) Porous sponges facilitate multifold improvement in transduction efficiency over a stagnant droplet (7x enhancement at a cell concentration of 20000 cells/µL, liquid volume of 50µL, and VCR of 1.65). (b) MicroCT and SEM images provide information on pore size distribution^25^, sponge surface, and the pore interconnection size distribution. Black squares show the mean value, and the error bars show standard deviation. (c) The liquid-porous interface position is experimentally measured using still images from a video of sponge liquid absorption. Liquid absorption into the sponge is computationally modeled as a 1D macroscale absorption. (d) The 1D liquid absorption model is calibrated using the experimentally measured interface position and then used to predict the absorption flow velocities experienced by the seeded liquid droplet. The time taken to absorb 200µL of liquid media is compared between simulation and experiment. (e) A 2D Periodic Representative Elementary Volume (REV) of the porous sponge is constructed to model microscale flow and particle trajectories (fluid - white pixels, solid - black pixels). (f) Microscale liquid flow is simulated inside the periodic REV to obtain the flow velocity distribution and (g) the probability distributions of the spatial fluctuation of streamwise (top) and transverse (bottom) flow velocity.

To deconstruct and quantify the physical contributions of dispersion and colocalization, we developed a multiscale simulation model. This framework was necessitated by the optical opacity of the sponges, which precludes direct visualization, and the prohibitive computational cost of full-scale numerical simulations across relevant length and time scales. Our prior experimental evidence identified interaction between convective flow and pore architecture as primary correlates of transduction efficiency, informing specific requirements for the model's physical fidelity. Specifically, the framework had to capture the causal hierarchy between these features: how capillary-driven absorption dictates bulk liquid velocity and how the stochastic pore architecture subsequently shapes this flow into tortuous microscale streamlines. It was essential to determine if this interplay between velocity and architecture generates a transport regime of sufficient magnitude to influence the statistics of cell-virus encounters in the sponges. By coupling these mechanisms through a reduced-order model, we sought to systematically quantify the physical forces at play and evaluate their individual contributions to the observed transduction enhancement.

#### Sponge liquid absorption induces microscale flow with stochastic spatial fluctuations

Liquid absorption into a macroporous sponge generates flow through a highly irregular pore network, where microscale velocity fluctuations are expected to emerge from stochastic geometry. To ground our transport model in physical reality, we first characterized the geometry and bulk absorption behavior of the macroporous alginate sponges. Using data from a previous X-ray microCT study and the open-source software OpenPNM, we quantified the pore size distribution, finding a mean pore size of 136 μm (**figure 3b**)^25^. Scanning Electron Microscopy (SEM) further revealed the degree of interconnectivity between adjacent pores (**supp. figure 5**), establishing the geometric constraints governing fluid flow. To describe the bulk liquid absorption, we developed a one-dimensional macroscale absorption model that balances capillary action, drag, and gravity. By tracking the liquid interface position from video recordings (**figure 3c**) and measuring liquid-air-sponge contact angle (**supp. figure 1**), we calibrated this model using MCMC to estimate the sponge's permeability (*k*_*c*_) as 0.0051 µm^2^ (95% CI [0.0044,0.0058]). The ability of this model to fit experimental data using a single constant permeability supports the assumption that the sponge remains hydrodynamically unchanged during absorption, justifying the neglect of poro-elastic effects. This macroscale model provides the time-dependent droplet position and absorption velocity (**figure 3d**), defining bulk liquid absorption dynamics that drive microscale flow within the sponge.

To resolve the microscale flow field responsible for particle dispersion, we next examined the fluid dynamics within the sponge pore network. We constructed a two-dimensional Representative Elementary Volume (REV) of the sponge's porous geometry by tightly packing circular pores^41^ with pore diameters drawn from the microCT-measured pore size distributions. The REV was rendered periodic, and the pore overlap was tuned to match the experimentally measured interconnection size (**figure 3b&e**). Although this 2D representation simplifies the full-scale 3D geometry, it statistically preserves pore size and interconnectivity – the dominant features governing hydrodynamic dispersion in creeping flow^42^. Using computational fluid dynamics, we simulated liquid flow through this 2D REV at the range of macroscale velocities predicted by our absorption model (*u*_*m*_ = 0.5, 1.0, 5.0, 12.5, and 25 µm/s). The simulations revealed that stochastic pore geometry induces significant spatial fluctuations in microscale velocity, producing tortuous streamlines (**figure 3f**). Analysis of the spatial distribution of the velocity fluctuations showed larger variance in the streamwise direction than the transverse direction (**figure 3g**). Furthermore, the streamwise distribution was skewed towards positive velocities, whereas the transverse distribution was symmetric and centered at zero. In both directions, the most probable velocity fluctuation corresponded to the zero-velocity, reflecting the dominance of no-slip boundary conditions at the pore walls and the large surface-area-to-volume ratio of porous materials. Since the flow occurs at very low Reynolds number (Re = 0.00007-0.004), the velocity fluctuations scale linearly with mean flow, rendering the nondimensional probability distributions invariant across flow speeds.

#### Cell and virus particles experience macroscale dispersion inside the pores of the sponge due to convective liquid flow

Given that the sponge pores induce stochastic velocity fluctuations, we next investigated whether these fluctuations generate hydrodynamic dispersion of the particles, providing cells and viruses with macroscale mobility beyond what is achievable with Brownian diffusion alone. We simulated the trajectories of more than 1,000 cell and virus particles randomly initialized within the 2D REV, assuming one-way coupling with the fluid flow, across a range of macroscale velocities predicted by the absorption model (*u*_*m*_ = 0.5, 1.0, 5.0,12.5, and 25 µm/s). To ensure the statistical robustness and independence from any specific geometry or initialization, results were averaged over five independent REV realizations (**supp. figure 6**) and five distinct initial particle distributions. The simulations revealed highly tortuous trajectories for both cells and viruses, reflecting the underlying flow streamlines, but with critical differences between the two particle types (**figure 4a**). As microscale inertial particles, cells frequently experience temporary immobilization upon impaction with pore walls^43,44^. In contrast, nanoscale viruses exhibit substantial Brownian motion, enabling them to migrate across flow streamlines. As a result, even initially co-located cell and viruses rapidly separate as they traverse the pore network, highlighting dispersion as a dominant transport mechanism within the sponge.

**Figure 4.**
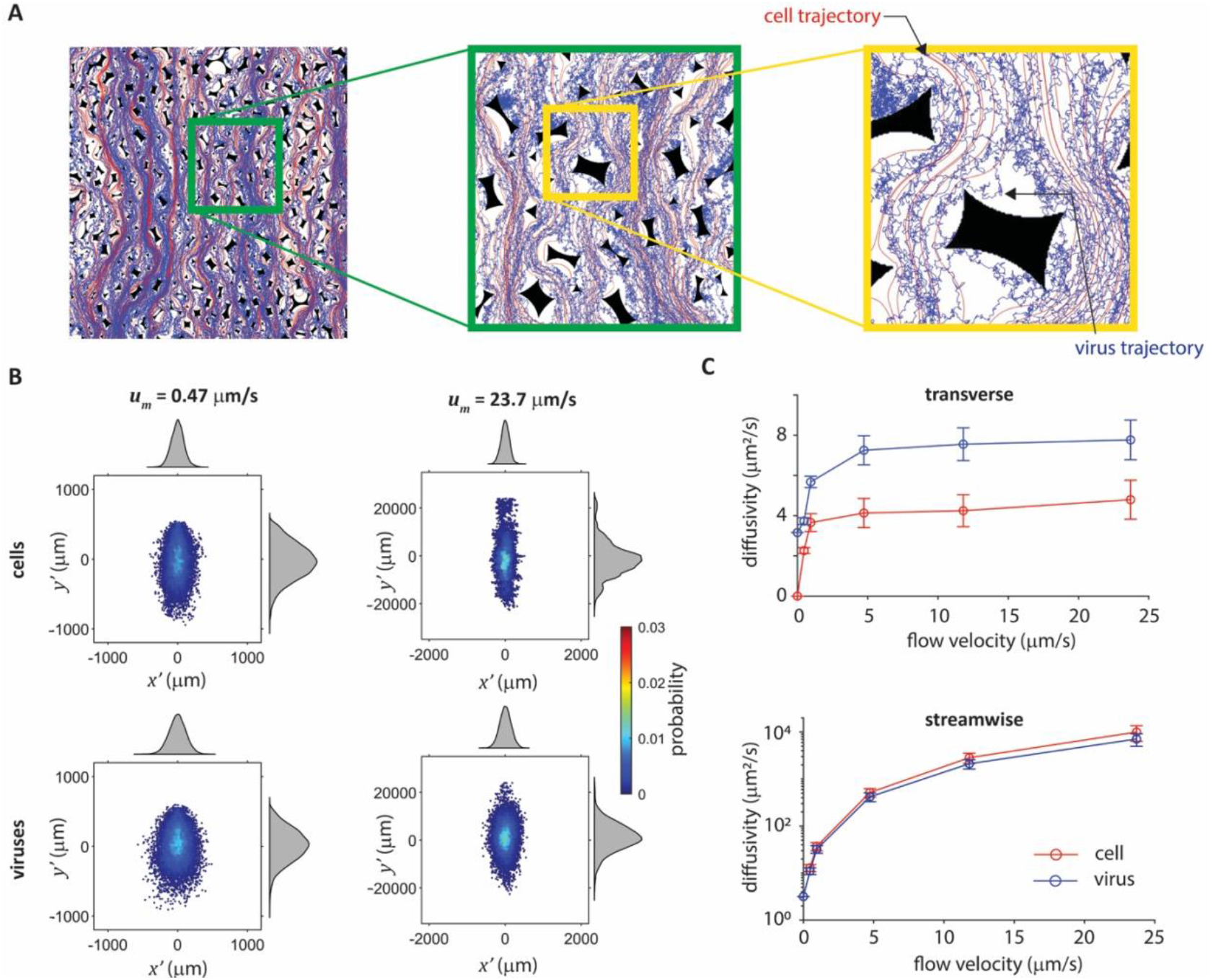
Microscale liquid flow particle trajectories are simulated inside the periodic REV. (a) Particle trajectories are simulated for both cells (red) and viruses (blue). Particle trajectories shown in this panel are for a flow velocity of 5µm/s. (b) Particle trajectories exhibit a fluctuating displacement from the mean macroscale flow trajectory due to dispersion in porous media. (c) Effective diffusivities are calculated for the cells and viruses at different liquid flow velocities.

To quantify the particle diffusion, we calculated the stochastic displacement (*x*_*p*_*'*) of each particle from the mean macroscale flow.

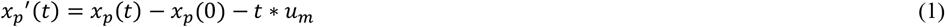

where *x*_*p*_(t) is the instantaneous particle position, *u*_*m*_ is the macroscale flow velocity, and *t* is the time interval. This expression follows directly from the particle equations of motion by assuming that the particle velocity is equal to the flow velocity due to the low Stokes number (*Re* < 0.004, *Stk* < 5×10^-7^), based on the advection-dispersion description of particle transport^45^. Analysis of particle positions revealed wide displacement distributions for both cells and viruses, with the viruses exhibiting greater dispersion due to the additional contribution from Brownian motion (**figure 4b**). Notably, the displacements were anisotropic, with substantially greater dispersion in the streamwise direction than the transverse direction, consistent with prior observations in porous flow^46^ and reflecting the higher variance in the streamwise fluid velocity fluctuations (**figure 3g**). This anisotropy intensified at higher flow velocities; as particles traverse more pores per unit time, compounding the dispersive effects of microscale velocity fluctuations. Such stochastic deviations from the mean flow trajectory is a hallmark of hydrodynamic dispersion in porous media^47^. Because stochastically distributed pore networks generate a greater degree of particle dispersion than uniform porosities^48,49^, the high heterogeneity of pore size in our sponges is expected to induce substantial particle dispersion that strongly influences cell and virus dynamics. This dispersion arises from a combination of streamline tortuosity^50–52^ and mass transport through pores of varying sizes and interconnectivity^53,54^, both of which drive particles away from the bulk flow path. We modeled this dispersion process by calculating an effective diffusivity (*D*_*eff*_) from the ensemble-averaged mean squared displacement of the particles over time, based on the Einstein diffusion model.

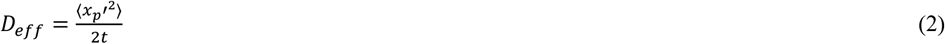

where ⦑ ⦒ denotes an ensemble average over all the particles. This procedure yielded a distinct effective diffusivity for cells and viruses at each macroscale flow velocity (**figure 4c, supp. table 2**). Notably, the longitudinal effective diffusivity in the streamwise direction exhibited a power-law dependence on the Peclet number (*Pe* = *u*_*m*_*d*_*pore*_/*D*_*mol*_), consistent with theoretical predictions for porous media in the range of 1 < *Pe* < 1000^55,56^. We then curve-fit these relationships (**supp. figure 7**) to develop a macroscale effective diffusion model. Interestingly, we observed that the particle displacements for both cells and viruses across different flow velocities followed a normal distribution (**figure 4b**). This result is well-supported by the Central Limit Theorem: although cells and viruses are advected by a velocity field that is not normally distributed, the mean displacement experienced by the numerous particles converges to a standard normal distribution. This observation enables efficient incorporation of the complex effects of microscale transport into our final macroscale model of transduction by assuming that the dynamics of the particles can be modeled as Gaussian noise whose spectral intensity is determined by the effective diffusivity at a given flow velocity.

### 2.3 Analysis of viral cellular transduction in a porous sponge

#### More cell-virus collisions occur in porous sponges resulting in improved transduction efficiency compared to a stagnant droplet

Having quantified the dispersive effects induced by sponge porosity, we next asked whether the magnitude of dispersion observed in the sponges, along with post-absorption pore colocalization, is sufficient to explain the experimentally observed enhancement in transduction. To address this question, we developed a comprehensive simulation model of viral transduction in porous sponges that integrates all the relevant physical mechanisms: the 1D liquid absorption model, the macroscale particle dispersion model, gravitational settling, and post-absorption particle colocalization within the sponge pores. As in the stagnant droplet simulations, the sponge geometry consisted of a periodically repeating representative sub-volume in the lateral directions (**figure 5a**). During liquid absorption into the sponge, the effective diffusivities of cells and viruses within the porous region are continuously updated based on the droplet's instantaneous absorption velocity. Once liquid absorption is complete, we assume that the cells and viruses become colocalized within the sponge pores, preventing further gravitational settling to the bottom of the entire liquid volume. Instead, particles become confined within a cubical volume equivalent to the mean pore size (136 µm). This assumption is based on experimental observations of cells visualized at the top of the sponge post-absorption^25^.

**Figure 5.**
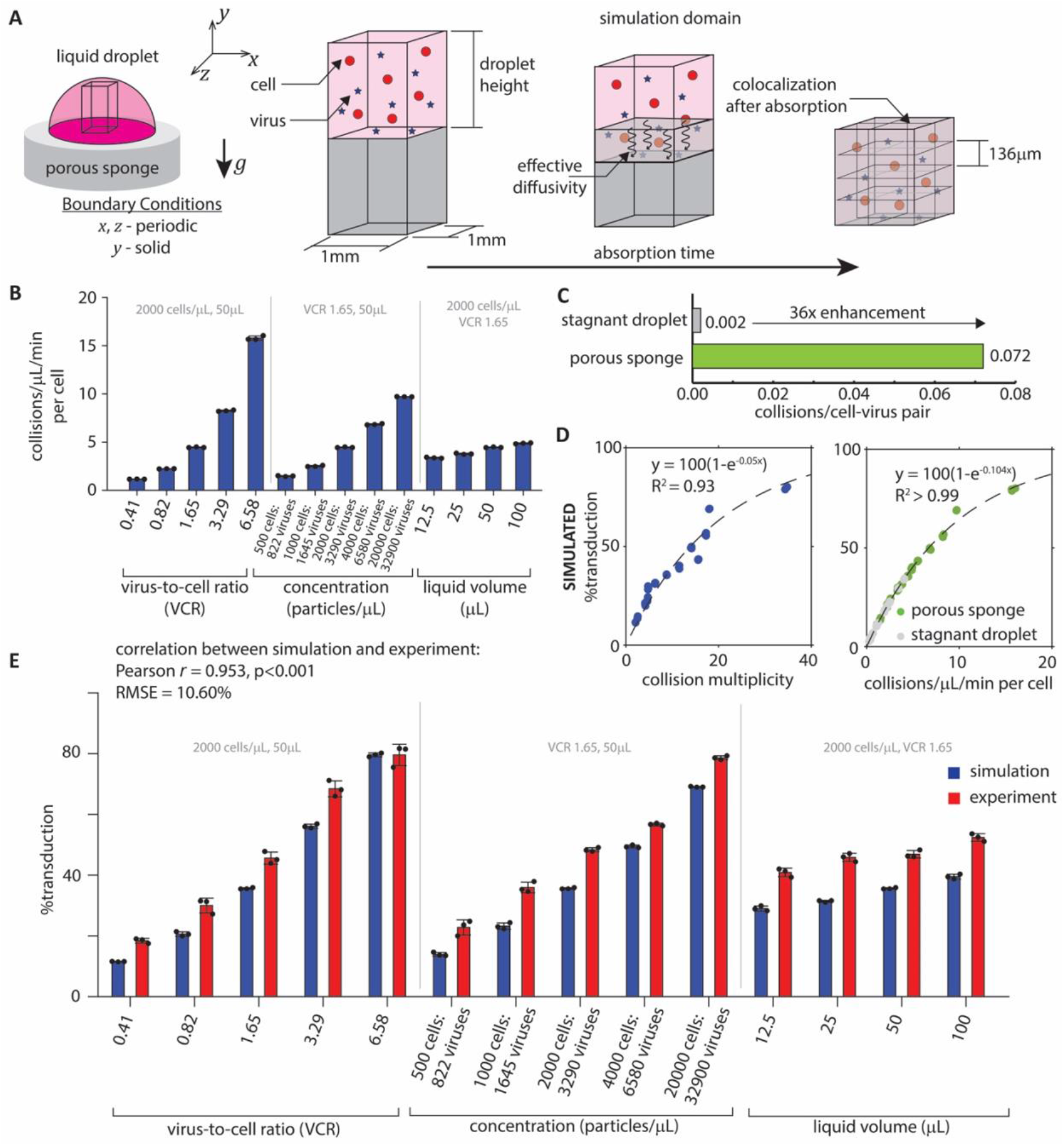
(a) Simulation geometry and boundary conditions used to model transduction in a porous sponge. (b) The simulation model predicts frequent cell-virus collisions occurring over time in the porous sponge. The constant parameters in each single parameter variation are a concentration of 2,000 cells/µL, VCR of 1.65, and liquid volume of 50µL. (c) Simulated cell-virus collisions per cell-virus pair per 1 µL volume across all tested values of VCR, cell-virus concentration, and liquid volume are substantially higher in the porous sponge when compared to the stagnant droplet. (d) Simulated transduction efficiency is strongly correlated to collision frequency and collision multiplicity. (e) Efficient transduction is predicted by the simulation and observed experimentally in porous sponges for various seeded liquid volumes, VCRs, and cell-virus concentrations.

Applying this integrated model, we predicted a substantially higher frequency of cell-virus collisions with liquid absorption into porous sponges compared to stagnant liquid droplets (**figure 5b**). Because sponge-mediated transduction is so efficient, we were forced to reduce the cell–virus concentration tenfold and the virus-to-cell ratio by half in the sponge simulations to avoid saturation and preserve sensitivity to parametric effects. Remarkably, even under these deliberately conservative conditions, this collision frequency in porous sponges remained higher than in stagnant droplets. By scaling the collision rate relative to the total number of cell-virus pairs, the model collapses these complex parameter dependencies into a single characteristic metric: a universal 36-fold increase in collision potential within sponges compared to stagnant droplets (R^2^>0.99; **figure 5c**).

Using the same transduction probability calibrated from the stagnant droplet experiments (*p*_*TE*_ = 0.0023), the model predicted the multifold increase in transduction efficiency observed experimentally in sponges (**figure 5e, supp. table 3**). Across a broad range of VCRs, cell-virus concentrations, and liquid volumes, the simulated transduction efficiencies showed strong agreement with experimental measurements in both stagnant droplets and porous sponges, supporting the model's predictive capacity. Crucially, this enhanced transport regime does not come at the expense of cellular health. Cell viability in the porous sponges remained consistently high (70-90%), with the lowest values observed at the extreme ends of the cell counts (**supp. figure 8**). While the simulation systematically underpredicted transduction efficiency (RMSE = 10.6%), this model error was minimized (RMSE = 5.83%) at a slightly higher transduction probability (*p*_*TE*_ = 0.0039), a value that remains within the range of uncertainty of *p*_*TE*_ that was previously estimated by MCMC.

#### Transduction in porous sponges is enhanced by macroscale particle dispersion during liquid absorption

The strong agreement between simulation and experiment supports the prediction that the enhancement in transduction is driven by increased collision multiplicity (**figure 5d**). Direct comparison of sponge and stagnant droplet systems under identical initial conditions (20,000 cells/µL, VCR 1.65, 50µL) highlights the dramatic contrast in the particle kinetics and spatial distribution within the sponge (**figure 6a**), enhancing the potential for cell-virus interactions. To identify the factors that contribute most to this enhancement, we analyzed the cell-virus collision history over the course of the experiment. The transduction process can be divided into two stages: a dynamic “absorption phase” and a subsequent static “colocalization phase". During absorption, particles experience enhanced effective diffusivity, while in the static phase they are colocalized within their immediate pore. Together, these mechanisms directly counter the dominant failure modes of the stagnant droplet system – slow Brownian diffusion and gravity-driven separation. Effective diffusivity during the dynamic phase actively mixes particles, sustaining access to new viral encounters and suppressing large-scale gravitational separation. In contrast, colocalization maintains a high local VCR and provides a three-dimensional interaction interface.

**Figure 6.**
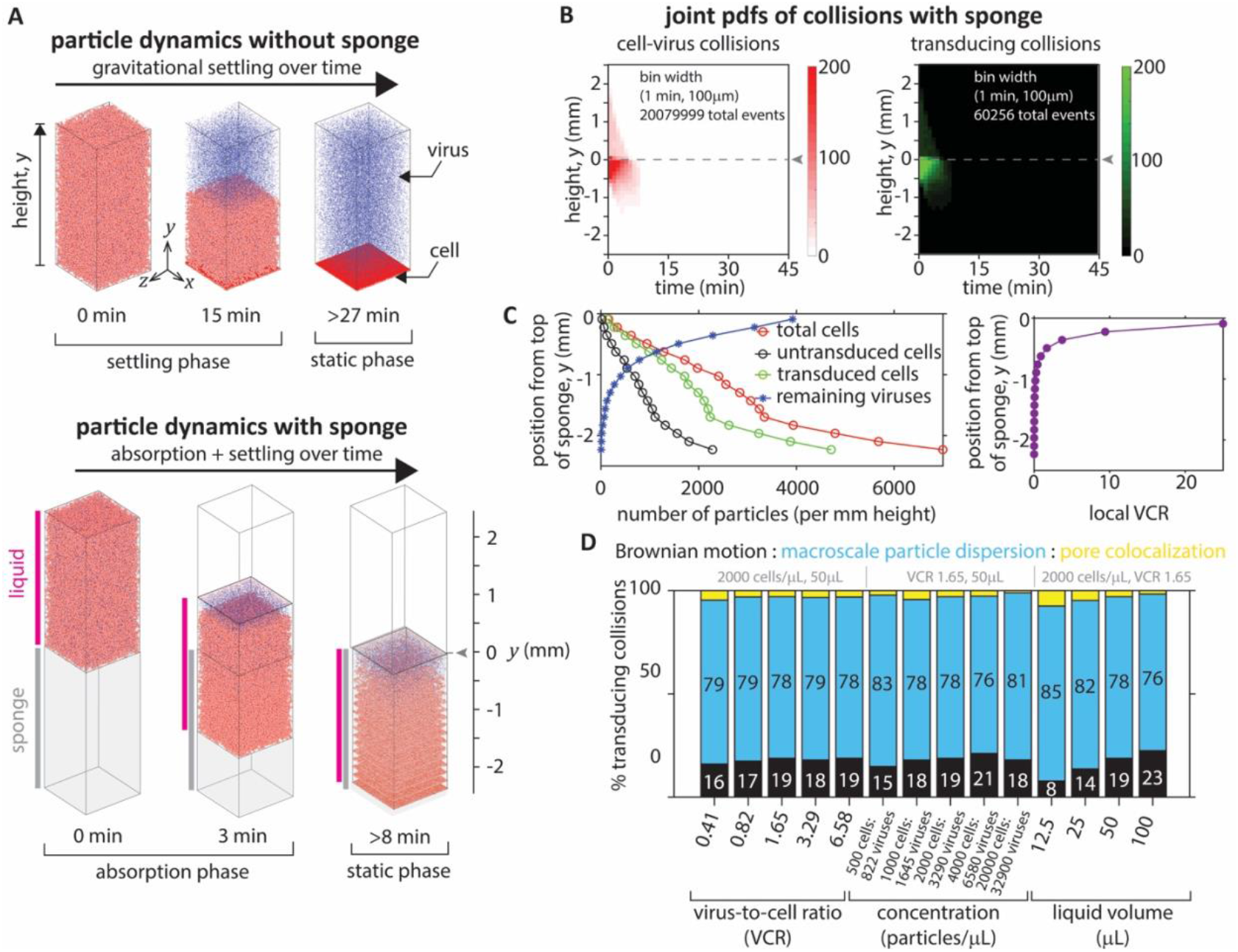
(a) Distributions of the cell and viral particles plotted over time show liquid absorption and gravitational settling. Visualizing cell settling with sponges shows that gravitational separation is not as drastic as with stagnant droplets with cells distributed throughout the sponge porosity even in the static phase. (b) Probability densities of both cell-virus collisions and successful transduction are greater during the dynamic absorption phase (*t* < 8 min). (c) The distributions of transduced cells, untransduced cells, and remaining viral vectors at the end of sponge transduction (45 min). Plots (a)-(c) are made for a cell-virus concentration of 20000 cells/µL, liquid volume of 50µL, and VCR = 1.65. (d) Fractional contributions of the different physical mechanisms towards transduction in porous sponges show macroscale particle dispersion is most significant. The constant parameters in each single parameter variation are a concentration of 2,000 cells/µL, VCR of 1.65, and liquid volume of 50µL.

Analysis of the collision histories revealed that the majority of collisions occur during the absorption phase (**figure 6b**). Collision rates peak early in absorption, coinciding with the highest liquid velocities (**figure 3d**) and decline as absorption slows down due to viscous drag. Additional collisions also occur in the liquid volume above the porous sponge due to Brownian motion during early settling. After absorption is complete, colocalization prevents gravitational settling and thereby preserves favorable local VCR. By categorizing each successful transduction event to its dominant physical origin, we quantified the contribution of each mechanism by calculating the number of transducing collisions that occurred due to it, assuming that collisions in each mechanism had the same likelihood of successful transduction (**figure 6d**). Macroscale dispersion during absorption accounted for the majority of transductions (72-85%). Brownian diffusion above the porous sponge accounted for between 8% and 23% of transductions, depending on absorption duration. The dominance of the dispersion mechanism is clear. This process physically displaces particles across multiple pore lengths (**figure 4a&b**), making it the most potent driver of transduction improvement.

Strikingly, only a small fraction of transductions (<10%) arose from pore-scale colocalization. This contrasts with a theoretical simulation isolating the mechanism of pore-scale colocalization that showed that colocalization improved transduction outcome, especially over prolonged time periods (**supp. figure 9**). This discrepancy is explained by considering the temporal sequence during transduction. During early stages, viral concentrations are highest and collisions are driven by Brownian motion. As absorption proceeds, macroscale dispersion drives a large number of productive collisions, depleting free viral particles. By the time absorption ceases and the colocalization stage begins, the viral concentration has already been substantially reduced. Consistent with this, analysis of cell and virus distributions at the end of the colocalization phase (**figure 6c**) shows that viruses are rarely co-localized with non-transduced cells, resulting in low local VCR and minimal additional transduction.

#### Optimizing transduction in porous sponges can improve virus utilization and control copy number

Viral vectors are a major cost and capacity bottleneck in CAR-T manufacturing because producing clinical-grade retroviral or lentiviral stocks is expensive, time-consuming, and constrained by GMP manufacturing capacity, contributing substantially to overall therapy cost and supply limitations^57,58^. These constraints motivate the development of novel transduction strategies together with quantitative models that relate transduction outcomes to viral vector utilization and vector copy number (VCN). Such models enable systematic optimization of both manufacturing efficiency and safety-relevant parameters. Consistent with these clinical and manufacturing needs, the combined model and experimental results demonstrate a sevenfold increase in the number of transduced cells in the sponge compared to a stagnant droplet under identical conditions (**supp. figure 10a**). The model reveals that this enhancement arises primarily from more frequent cell-virus binding, with many cells binding more than one virus (**supp. figure 10b&c**).

By analyzing the histogram of bound viruses per cell, we quantified the efficiency of viral vector utilization in both the stagnant droplet and sponge systems. We found that although increasing either the VCR or the cell-virus concentration boosts transduction, increasing cell-virus concentration is far more effective at utilizing the available viruses and minimizing waste. At high concentrations, the sponge utilized up to 90% of the initial virus, compared to a maximum of °15% in the stagnant droplet (**supp. figure 10d**). We derived an analytical estimate for virus utilization by fitting a Poisson probability distribution to the simulated result of bound viruses per cell and observed that virus utilization at a given VCR is determined by transduction efficiency (**supp. figure 10e**). These findings suggest that optimizing for particle concentration, rather than VCR alone, is key to achieving maximum transduction efficiency while simultaneously reducing the volume of expensive viral vector required for manufacturing.

While the sponge system dramatically improves efficiency, our model reveals an inherent trade-off between transduction efficiency and the resulting vector copy number (VCN). Specifically, higher transduction efficiency is accompanied by more viruses binding per cell (**supp. figure 10f&g**), revealing a logarithmic relationship between transduction efficiency and the number of bound viruses per cell. Notably, the model-predicted relationship between bound viruses per cell and transduction efficiency parallels the experimentally reported correlation between transduction efficiency and VCN in CD34^+^ cells^59^. This implicit link between VCN and transduction efficiency creates an optimization challenge, as elevated VCN, while linked to therapeutic potency, must be carefully balanced against safety concerns^60–62^.

## 3 Discussion

Reactive biological processes that depend on rare encounters between particles of disparate size and mobility are fundamentally constrained by transport physics. Traditionally, the governing mechanisms of such encounter-limited systems have been relegated to reductive, well-mixed idealizations, closed by empirical models that lack robustness and remain inextricably tethered to specific geometries, methods, and protocols. This study fundamentally transforms this landscape and establishes a generalized physical framework that accounts for the intrinsic spatiotemporal heterogeneity of these processes. Applying this framework, we resolve the collision dynamics emergent when advection, geometry, and stochasticity interact across scales to provide a transport-level, physical explanation for the striking enhancement of cell-virus interactions in biomaterials.

By integrating experimental measurements with a multiscale simulation model, we demonstrate that viral-cellular transduction in stagnant liquid is inefficient because diffusion-dominated transport of nanoscale viruses cannot compensate for gravity-driven segregation of microscale cells. In contrast, absorption into a porous sponge drives the system into a fundamentally different transport regime. Capillary-driven flow through a stochastic pore network generates strong hydrodynamic dispersion, converting unidirectional advection into effective macroscale mixing. This dispersion dominates the encounter statistics, accounting for the majority of productive cell-virus collisions. Secondary contributions arise from Brownian diffusion both above the sponge and from geometric confinement following absorption, which provides a three-dimensional interface for cell-virus interaction.

The enhancement driven by macroscale particle dispersion persisted across all tested virus-to-cell ratios, particle concentrations, and liquid volumes, indicating that the effect is robust and geometry-mediated rather than parameter specific. This geometry-induced hydrodynamic dispersion reshapes collision statistics, converting diffusion- and segregation-limited dynamics into a regime dominated by sustained, high-multiplicity encounters. While pairwise recurrence remains fixed by molecular diffusivity, collision multiplicity is a transport-controlled quantity that can be dramatically amplified by dispersion in heterogeneous flow fields.

Crucially, the model reveals that this superior transport enables the sponge to utilize up to 90% of the initial viral dose – a 11-fold increase over the °8% utilization observed in stagnant droplets under the same conditions. Despite a sevenfold improvement in overall transduction efficiency in the sponge, the maximum observed number of bound viruses remains as low as 1.6 viruses per cell. This suggests that the sponge transduction mechanism achieves high efficiency and exceptional virus utilization while maintaining a favorable safety profile regarding the number of genomic integrations. Given that viral vector production is a major cost and capacity bottleneck in CAR T manufacturing, such high utilization rates are essential for overcoming the supply limitations that currently restrict the delivery of these life-saving treatments.

While our model shows strong agreement with experimental trends, it operates with certain simplifying assumptions that open clear avenues for future investigation. Our analysis identifies sources of uncertainty that limited model accuracy, primarily with the quantification of functional viral particles and calibration of model parameters. As a result, in porous sponges, the model underestimated transduction efficiency with a root mean square error of 10.6%. We determined that this error could be minimized to 5.83% by adjusting the transduction probability to a value that is within the 95% credible interval. Additional sources of uncertainty in transduction probability include the estimation of liquid absorption rate or the persistence of viral vectors within the sponge pores beyond the 45-minute transduction window. Whereas, in stagnant liquid, the effect of cell crowding during culture caused poor cell viability at high concentrations and the formation of multilayers of settled cells potentially led to restricted viral access in the static phase.

Future extensions incorporating direct numerical simulation on full-scale 3D tomographic reconstructions of the sponge, high spatiotemporal resolution characterization of liquid absorption, and precise control of viral dilution at the transduction endpoint could offer even greater fidelity and enhance the model's predictive accuracy. Furthermore, our experimentally calibrated transduction probability, while effective, assumes the outcome of an individual cell-virus collision is independent of the local fluid dynamics. Whether collisions in a high-velocity flow regime yield the same outcome as those in a static environment is a fundamental question that presents a significant technical challenge reserved for future experimental work.

By establishing a quantitative link between microscopic particle kinetics and macroscopic reaction outcomes, this study demonstrates that cell-nanoparticle interactions in encounter-limited systems can be fundamentally controlled by transport physics. By capturing these physical drivers within a stochastic collision framework, we move from empirical observation to mechanistic prediction of critical encounters in biomedicine, such as in viral-cellular transduction. This provides a physics-based paradigm for the in-silico design of porous media and protocols, enabling the systematic optimization of scalability and cost-effectiveness in cell therapy manufacturing.

## 4 Materials and methods

### 4.1 Experimental details

#### 4.1.1 Sponge Synthesis

Alginate sponges were prepared following our previous work^24,27,63^. Ultrapure alginate (Pronova, MVG) was dissolved in deionized water by stirring for at least one hour to create a 2% w/v solution before being mixed with an equal volume of 0.4% w/v of calcium gluconate solution for 15 min. The final solution has an alginate concentration of 1% w/v and calcium concentration of 0.2% w/v. The resulting mixture was cast 1 mL per well in a 24-well non-coated tissue culture plate and frozen at -20 °C overnight. The frozen sponges were lyophilized for 72 hours before being removed, vacuum sealed with desiccants and stored at 4 °C until used.

#### 4.1.2 Scanning Electron Microscopy

The sponge porosity was analyzed on a Hitachi SU3900 variable pressure scanning electron microscope. Samples were analyzed at 70 Pa with a nitrogen backfill gas. Images were captured using the ultra-variable detector with a 20kV accelerating voltage. The sponge image analysis was performed on the top surface to measure the size of the pore interconnections using ImageJ/Fiji. Sponge pore interconnections were differentiated from the pores based on the orientation of the void with respect to the sponge wall. Voids aligned with the sponge walls were assumed to be interconnections. Voids orthogonal to the sponge walls were assumed to be pores. Sponge pore size distribution used in this study is taken from a previous study that analyzed sponge porosity with X-ray micro-CT image analysis^25^.

#### 4.1.3 Contact angle and liquid absorption imaging

Still photographs and videos were acquired using a mobile phone camera mounted on the workbench. Still images were acquired at a resolution of 12MP, and videos were acquired at 1920×1080 pixels and 60 fps. To measure the contact angle between the sponge, liquid media, and air, the sponge was first compressed into the shape of a flat disk to prevent liquid absorption. Complete cell culture media was pipetted onto the flattened sponge, and the droplet shape was photographed. The same procedure was repeated to measure the contact angle at the bottom of a non-treated 24-well plate. To measure the liquid absorption into the sponge, 275µL of complete cell culture media was pipetted onto the top of the sponge to cover the entire surface area and allowed to absorb into the sponge. The absorption was captured by video and then analyzed frame-by-frame to determine the interface position. The sponge height was measured prior to absorption as 4.75 mm and used to calibrate the image pixels to a physical length scale in the video frame.

#### 4.1.4 Liquid medium density, cell material density, and cell diameter measurement

The material density of the complete cell culture media was measured as 1007.753 ± 2.466 kg/m^3^ (*n* = 10) by repeatedly weighing a 1mL volume at room temperature (294 K). The dynamic viscosity of the cell culture media is assumed to be 0.000958 Pa·s based on the observed value for RPMI-1640 with 10%v/v fetal bovine serum^64^. To measure the cell material density and diameter, brightfield z-stack images of Jurkat cells settling in the liquid medium were acquired on a Nikon Ti2 confocal microscope equipped with a fast-scanning piezo stage and 20X, 0.8 Numerical Aperture Objective (Nikon MRD70270). 1 million Jurkat cells were suspended in 500µL of complete cell culture media and pipetted onto an optical-bottom 24-well plate. A sub-volume of the cell suspension at the bottom of the well (665.6 × 664.3 × 180 µm; 2048 × 2044 × 36 pixels) was imaged at 2.76s per volume for 44 repetitions (121s). The cell settling process was repeated 6 separate times and each time starting from a thoroughly mixed suspension. Cell settling velocity was calculated by manually tracking cells over time in ImageJ/Fiji, recording their initial and final positions along the direction of gravity, and then calculating the settling velocity. Cells were chosen for tracking across the entire imaging volume. The diameter of each cell was calculated as 10.60 ± 0.86 µm (*n* > 100) by drawing a circular mask around the cell to measure the cell area, assuming a circular cross-section. Cell material density was calculated as 1029.056 ± 6.193 kg/m^3^ (*n* > 100), from the settling velocity and cell diameter by applying Stokes' law and assuming that the settling velocity is equal to the terminal velocity. Virus material density is also assumed to be 1029 kg/m^3^, which is similar to reported ranges^65,66^, considering that virus weight is negligible when compared to its Brownian force.

#### 4.1.5 Virus production and functional titer

GFP γ-retrovirus was produced using a stably transfected FLYRD118 packaging cell line. The physical titer of the viral stock was quantified using Nanoparticle Tracking Analysis (NTA) with a ParticleMetrix Zetaview instrument. Samples were first diluted in sterile, particle-free phosphate-buffered saline (PBS) to fall within the instrument's optimal concentration range for measurement. The concentration (2.2 × 10^10^ physical viral particles/mL) and size distribution (150 ± 41nm) were then determined by the instrument's software, which calculates these parameters based on the Stokes-Einstein diffusivity model. The functional viral titer was determined using a standard flow-cytometric assay. In this approach, the functional titer was determined by transducing 1 million Jurkat cells in triplicate using 48-well alginate sponges with 50 µL of respective dilutions (no dilution, 5x, 20x, 80x, and 320x) of concentrated virus. Dilutions that result in 10–40% GFP+ cells were used to calculate the viral functional titer using the following equation and the resulting titers were averaged: titer (TU mL^−1^) = (cell number used for infection × percentage of GFP+ cells)/(virus volume used for infection in each well × dilution fold). This functional titer value (2.07 × 10^7^ TUs/mL) is then used to prepare the experimental cases, and the cells are analyzed to determine experimental transduction efficiency. However, a more robust quantitative assessment of the functional titer is applied to set up the simulation model and calculate the simulated transduction efficiency as detailed in supplementary appendix A. In this approach, we applied probabilistic modeling to estimate the posterior probability distribution of the functional viral titer using Markov Chain Monte Carlo and then assumed the median value as the functional titer (3.40 × 10^7^ TUs/mL). Following this approach, we ensured that the functional titer is rigorously estimated considering different cell and viral concentrations and different transduction methods (sponge and droplet) over the entire gamut of observed transduction efficiencies (0-100%) with root mean squared error less than 6%. The functional viral titer was estimated using a separate, mechanism-agnostic probabilistic model. This ensured that the numerically estimated titer was an independent input that did not bias the simulation model.

#### 4.1.6 Transduction protocol

Transduction experiments were performed using γ-retroviral vector encoding for GFP produced in house. The GFP retroviral supernatant was concentrated twentyfold using Amicon centrifugation filters (MWCO 100 kDa, Millipore) at 1500g for 10 min in a swinging bucket rotor. Concentrated retrovirus and complete cell culture media were used to resuspend a prepared pellet of Jurkat cells in the desired cell-virus concentration and liquid volume. This suspension was immediately pipetted onto either the top of the dry macroporous alginate sponges or directly at the bottom of an individual well in a 24-well plate (stagnant liquid droplet). A no virus control was prepared by resuspending 1 million Jurkat cells in 50µL complete cell culture media and adding to the bottom of an individual well. The cell-virus suspension was incubated at 37 °C for 45 min, after which 2 mL of complete cell culture media (RPMI 1640 with L-glutamine (Gibco), supplemented with 10% w/v of fetal bovine serum (Corning) and 100 U mL^−1^ penicillin (Gibco), and 100 µg mL^−1^ streptomycin (Gibco)) was added to each well. After 72 h of incubation, sponges were dissolved with 1 mL of 0.05 M EDTA to isolate cells. In both sponge and stagnant liquid droplet groups, the cells were washed twice with DPBS, stained with anti-human anti-CD3 antibody (BD Biosciences, APC-Cy7) and live/dead stain (Sytox Blue, Fisher Scientific) according to manufacturer's protocol, filtered and prepared for flow cytometry. All transduction experiments were performed on the same day using the same stock of Jurkat cells and GFP γ-retrovirus, and then subsequently analyzed for GFP expression after 72 h.

#### 4.1.7 Flow cytometry

All samples were analyzed on an Attune NxT (ThermoFisher) flow cytometer with a minimum of 5000 live CD3+ events acquired per sample. All events were gated on Jurkat cells by FSC, FSC singlets, viable cells, CD3+, and then GFP+. Transduction efficiency was calculated from the final GFP+ population. Analysis of data was performed using FlowJo following the gating strategy in **supp. figure 11**.

### 4.2 Numerical details

Expanded details of the governing equations and numerical method are provided in supplementary appendix B.

#### 4.2.1 Simulating transduction at the macroscale level

Liquid absorption into the porous sponges is simulated by using a 1D liquid absorption model to predict the position and the velocity of the liquid droplet over time by calculating the balance of capillary, drag, and gravitational forces. Cells and viruses are assumed to be hard spherical particles and their dynamics are simulated in 3D by using the discrete particle model (DPM) considering forces of hydrodynamic drag, Brownian motion, and gravity^67^. In sponge simulations, an additional “effective diffusivity” force is considered to model the dispersive mechanism of the sponge pores. Particles are assumed to elastically bounce back at both liquid-solid and liquid-air interfaces^31^. Jurkat cells are non-adherent and they do not stick to the solid walls made of alginate. Electrostatic attraction is negligible as both the ?-retroviruses and alginate sponge walls carry a net negative charge. Collisions between cell and virus particles are calculated when the distance between the centers of a cell and a virus is less than or equal to the sum of the radii. T cell doubling (>8ph^68–70^) and viral half-life (2-9 h^37–40^) are not considered in the model since their time scales are greater than the incubation time of the cells and viruses (45 min).

Transduction is predicted from the cell-virus collision history by using a probabilistic model which simulates a Bernoulli trial defined by transduction probability, *p*_*TE*_. To validate our approach, we employed a strict separation between model calibration and model prediction. The transduction probability was calibrated using a single subset of experimental data: the variation of particle concentration in stagnant droplets. This calibrated value was then held constant to predict transduction outcomes across all other experimental conditions, including variations in virus-to-cell ratio (VCR), liquid volume, and the entirely distinct physical regime of the porous sponge. Transduction probability (*p*_*TE*_) is numerically estimated from the experimental transduction efficiency by using Markov Chain Monte Carlo. The uncertainty in the simulation predictions of experimental outcome is quantified by estimating the posterior probability distribution of *p*_*TE*_, calculating the mean value and the 95% credible interval.

#### 4.2.2 Simulating particle dispersion at the microscale level

At the microscale level, fluid dynamics and particle motion are modeled in 2D. A representative geometry of the sponge pores is created by tightly packing circles by minimizing the distance between their centers^41^ to mimic the ice crystallization process during cryogelation. The diameters of the circles were randomly drawn from the probability density distribution of the sponge pore size distribution obtained from microCT image analysis. The circle packing is then padded with circles drawn from the sponge pore size distribution to make the representative simulation volume rectangular and fully periodic. The circles in the packing are then overlapped such that the mean and variance of interconnection size in the representative model are identical to that of the SEM measurements. While this 2D representation simplifies the full 3D geometry, it was constructed to statistically preserve the pore size distribution and interconnectivity measured via microCT and SEM. In the creeping flow regime (*Re*<0.004) relevant to this study, hydrodynamic dispersion is primarily governed by these pore-scale statistics rather than 3D inertial flow structures, making this 2D approximation a robust estimator of particle kinetics.

A computational fluid dynamics model was developed in ANSYS Fluent^71^ to calculate the microscale liquid flow velocity and pressure distributions inside the representative sponge pore geometry by solving the steady, incompressible Navier-Stokes equations and the pressure Poisson equation. Cells and viruses are simulated using a 2D DPM model^67^ considering forces of hydrodynamic drag, pressure gradient, Brownian motion, and gravity. Shear stress gradient, Saffman, virtual mass, and Basset forces were omitted since they were at least 3 orders of magnitude smaller than the drag force. The effect of hydrodynamic stress in deforming cells is not considered since the simulation predicted that the shear stress inside the sponge pores is 3 orders of magnitude smaller than the stress required for mechanoporation^23^. The simulated cell and virus trajectories are then independently analyzed to calculate the effective diffusivity of the particles at different volume-averaged flow velocities.

## Acknowledgements

This work was funded by grants R33CA281875, R01EB019409 and R37CA260223 from the National Institutes of Health and DMS-2451660 from the National Science Foundation. M. M. acknowledges fellowship funding from T32GM141887. M.M. and V.S. thank the North Carolina State University Comparative Medicine Institute for their support through the Student Ideation Award. We acknowledge the computing resources provided by North Carolina State University High Performance Computing Services Core Facility (RRID:SCR_022168) and the University of North Carolina at Chapel Hill Research Computing group. Flow Cytometry experiments were performed in the UNC Flow Cytometry Core Facility. The UNC Flow Cytometry Core Facility (RRID:SCR_019170) is supported in part by P30 CA016086 Cancer Center Core Support Grant to the UNC Lineberger Comprehensive Cancer Center. Research reported in this publication was supported in part by the North Carolina Biotech Center Institutional Support Grant 2017-IDG-1025 and by the National Institutes of Health 1UM2AI30836-01. The content is solely the responsibility of the authors and does not necessarily represent the official views of the National Institutes of Health. Scanning Electron Microscopy (SEM) was performed at the Analytical Instrumentation Facility (AIF) at North Carolina State University, which is supported by the State of North Carolina and the National Science Foundation (award number ECCS-2025064). The AIF is a member of the North Carolina Research Triangle Nanotechnology Network (RTNN), a site in the National Nanotechnology Coordinated Infrastructure (NNCI). We thank Chuck Mooney for training and support on the Hitachi SU3900 in the AIF. Nanoparticle Tracking Analysis was performed at the Nanomedicines Characterization Core Facility, Center of Nanotechnology in Drug Delivery, UNC School of Pharmacy. We thank Leo Kuo for his support with Nanoparticle Tracking Analysis. We thank Madelyn VanBlunk for sharing preliminary experimental data and insightful discussions during this project. We thank Israt Jahan Tulip for her support with sponge synthesis.

## Supplementary figures

**Supplementary figure 1.**
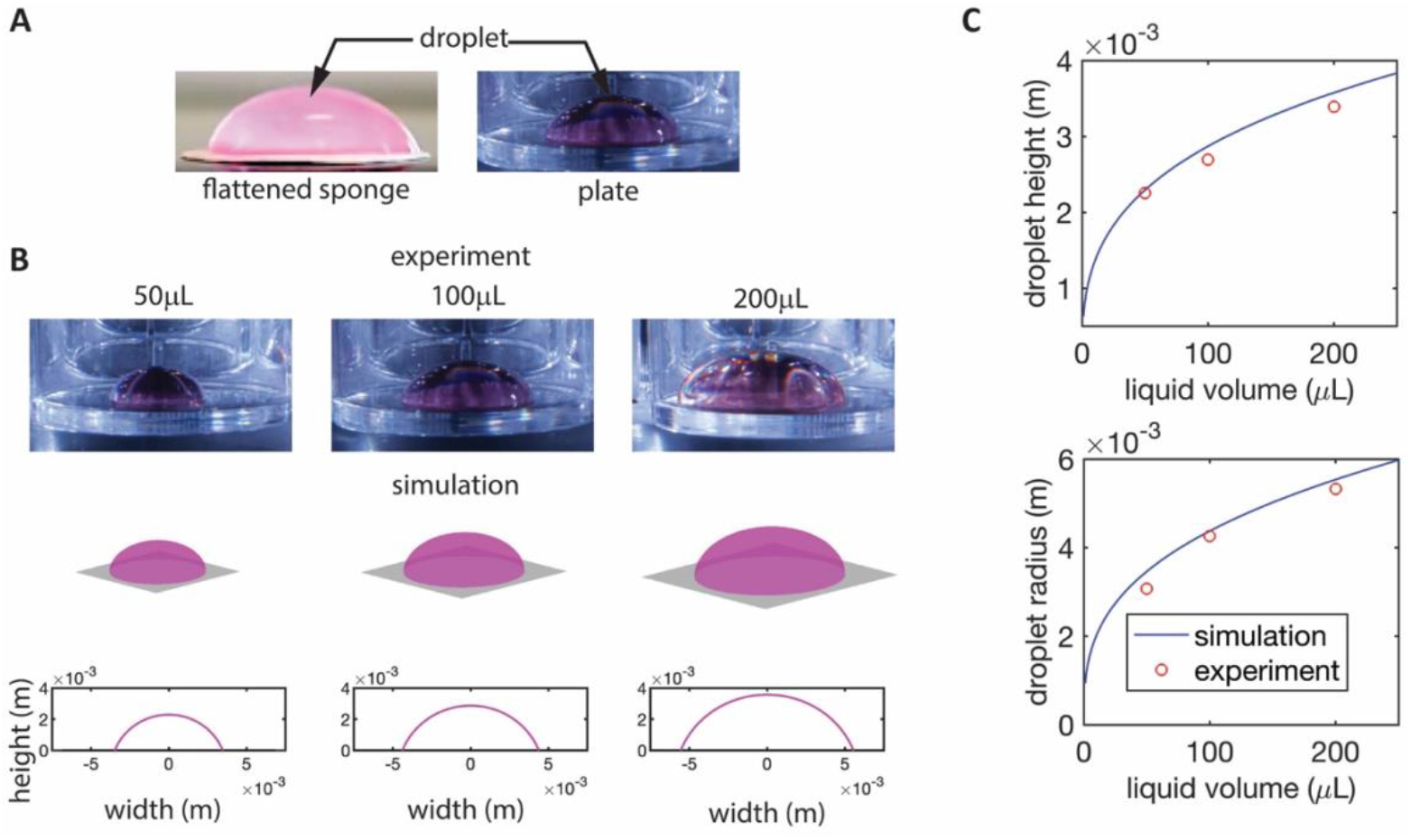
(a) The contact angle of the liquid droplet with both the flattened sponge disk and the well-plate materials are estimated as 68°. We measured the contact angle at the bottom of a non-treated 24-well plate. We measured the contact angle of the liquid-air interface at the sponge walls by seeding a droplet of the liquid medium on a sponge that had been compressed into a disc shape to remove the porosity. This prevented absorption and allowed the droplet to form a static shape on top of the calcium-crosslinked alginate material. (b) Droplet shape is simulated at different liquid volumes, and the resulting droplet height and radius is compared against the experimentally measured value (c). Droplet shape simulation model details are provided in supplementary appendix B.

**Supplementary figure 2.**
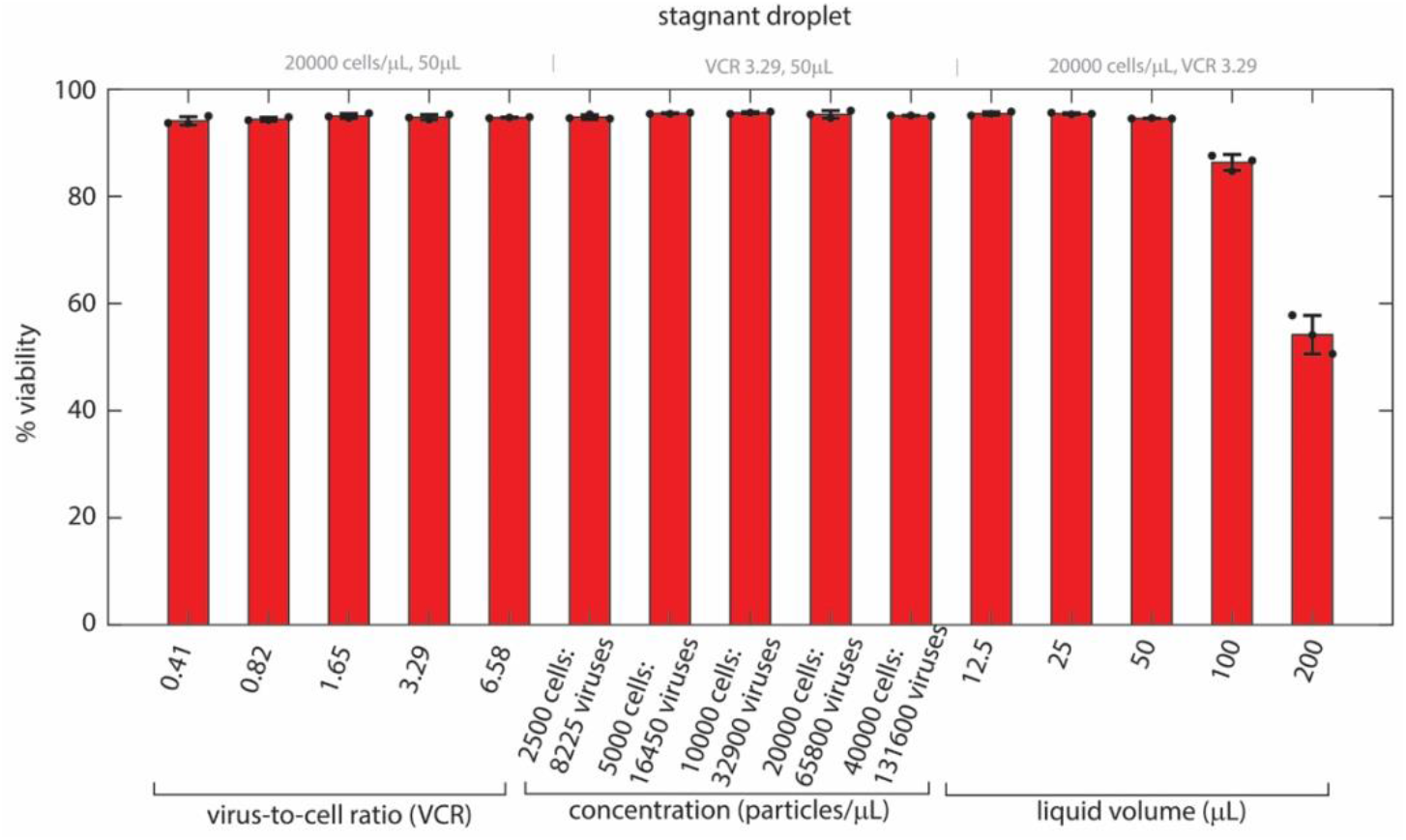
Experimentally measured cell viability in the stagnant liquid droplet for different VCRs, particle concentrations, and liquid volumes. Cell viability was analyzed after transduction protocols and 72h cell culture.

**Supplementary figure 3.**
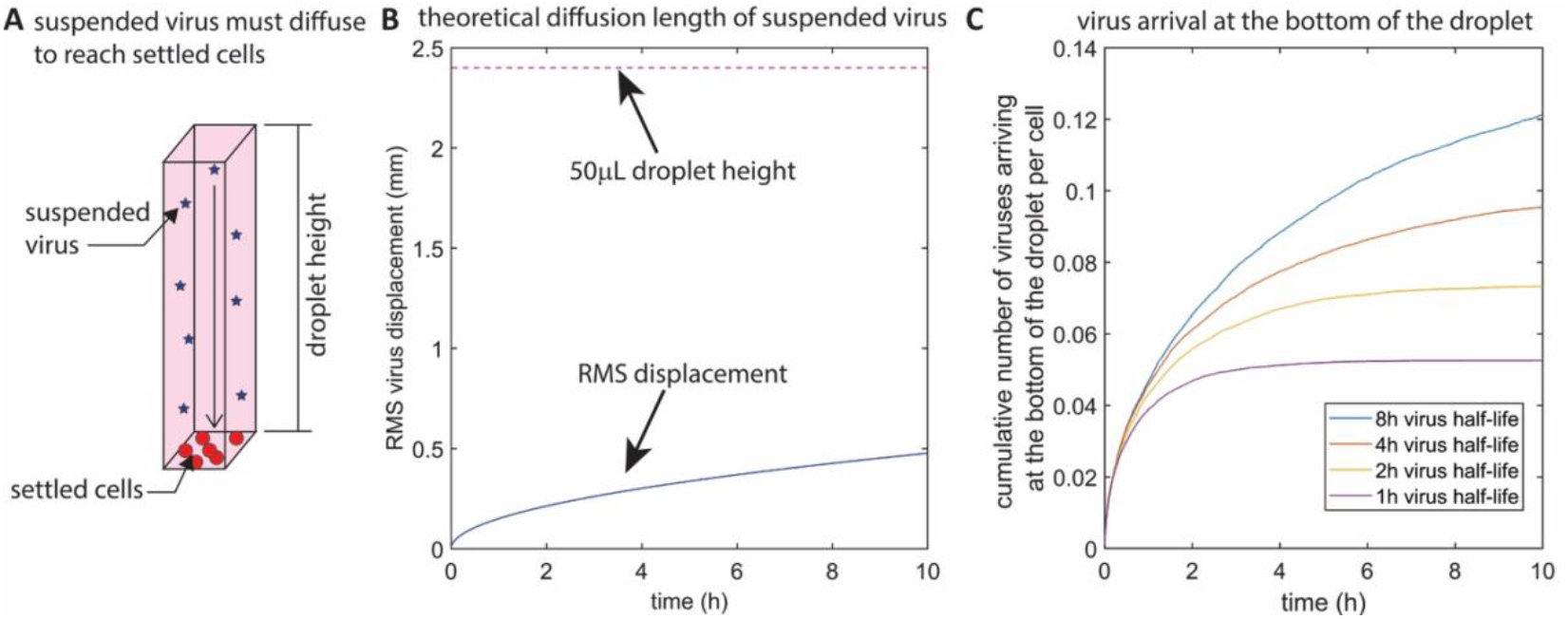
(a) When the cells settle to the bottom of the stagnant liquid droplet due to gravity, the suspended viruses must diffuse large distances to transduce the cells. (b) The root-mean-square (RMS) displacement of the virus (blue solid line) predicted by the Stokes-Einstein theoretical model indicates that viruses could only diffuse a root-mean-squared displacement of °0.5mm over a 10-hour period, which is only 20% of the droplet height (magenta dashed line). (c) During the 45 min incubation period used in the experiments, less than 15% of the total virus population reach the settled cells, and even fewer would bind to the cells due to low probabilities of collision and transduction. Factoring viral half-life due to degradation at 37°C, the cumulative number of viruses arriving at the settled cells is considerably low even at the 10-hour time point, where 1 virus is shared between 8 to 20 cells depending on the half-life of viral degradation. Calculations are made for a cell-virus concentration of 20000 cells/µL, liquid volume of 50µL, and VCR = 1.65.

**Supplementary figure 4.**
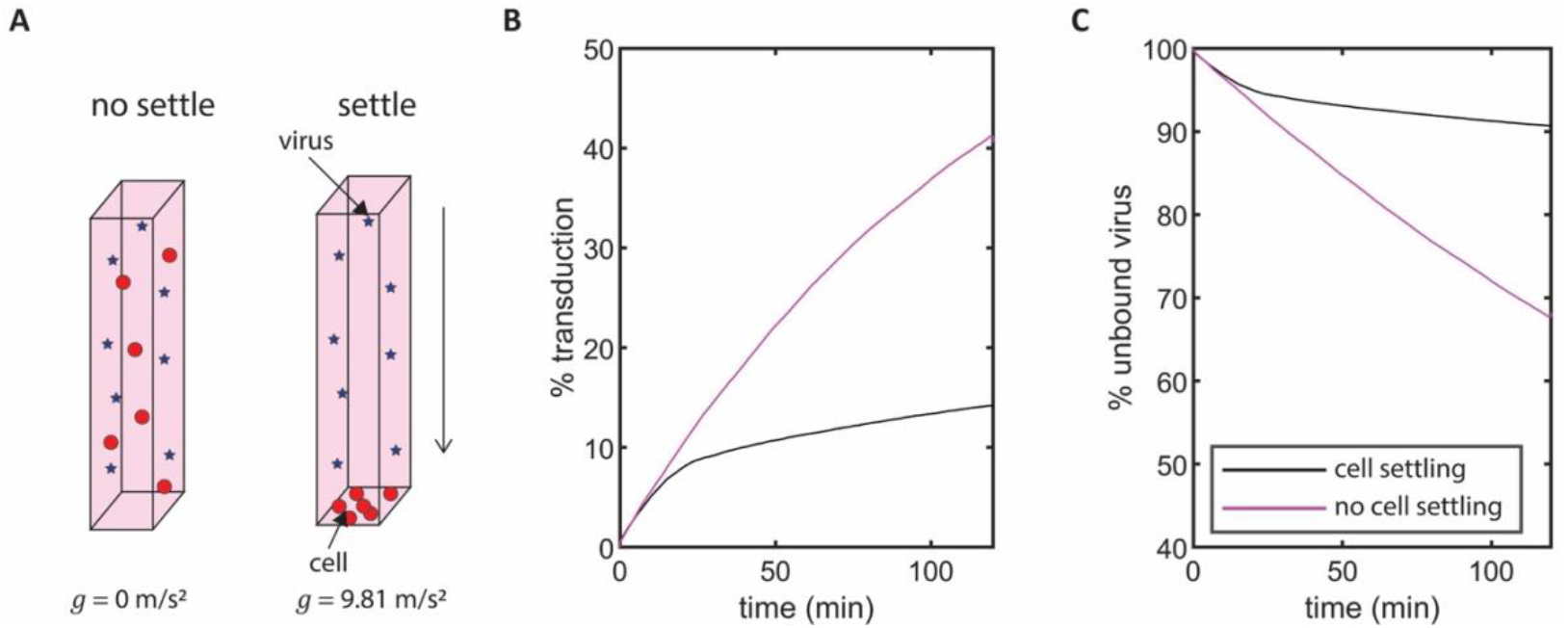
(a) Transduction is simulated for cases where gravitational settling either exists or does not exist by specifying the gravitational acceleration. (b) Transduction consistently increases over time when the cells are prevented from settling when compared to saturated transduction when the cells are allowed to settle. (c) When settling is prevented, more viruses are utilized by allowing the viruses to collide with the distributed cell population. This highlights that gravitational settling and not Brownian motion is responsible for poor transduction efficiency in stagnant liquid droplets. Calculations are made for a cell-virus concentration of 20000 cells/µL, liquid volume of 50µL, and VCR = 1.65.

**Supplementary figure 5.**
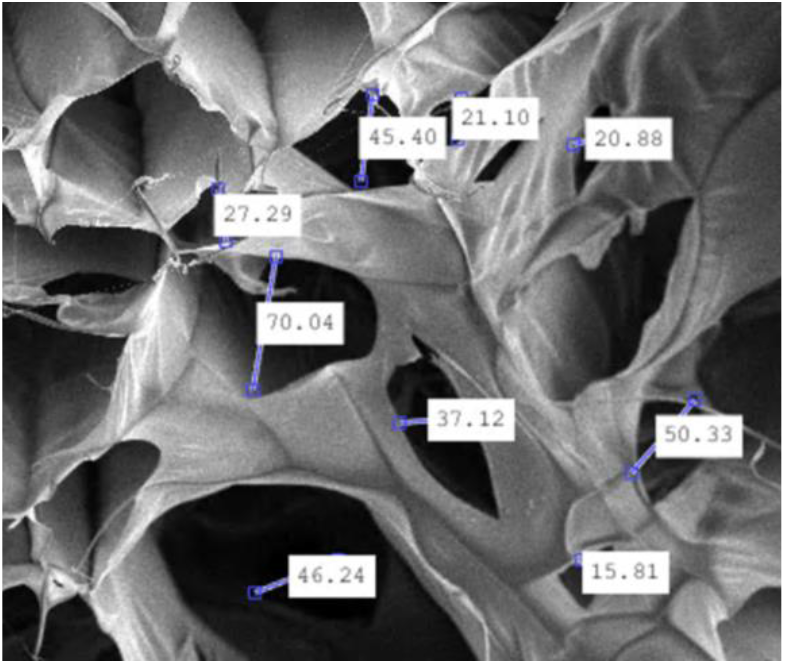
SEM image of the sponge surface porosity with pore-to-pore interconnections labeled and measured. The labels show the pore interconnection size in micrometers. A total of 20 interconnections were measured.

**Supplementary figure 6.**
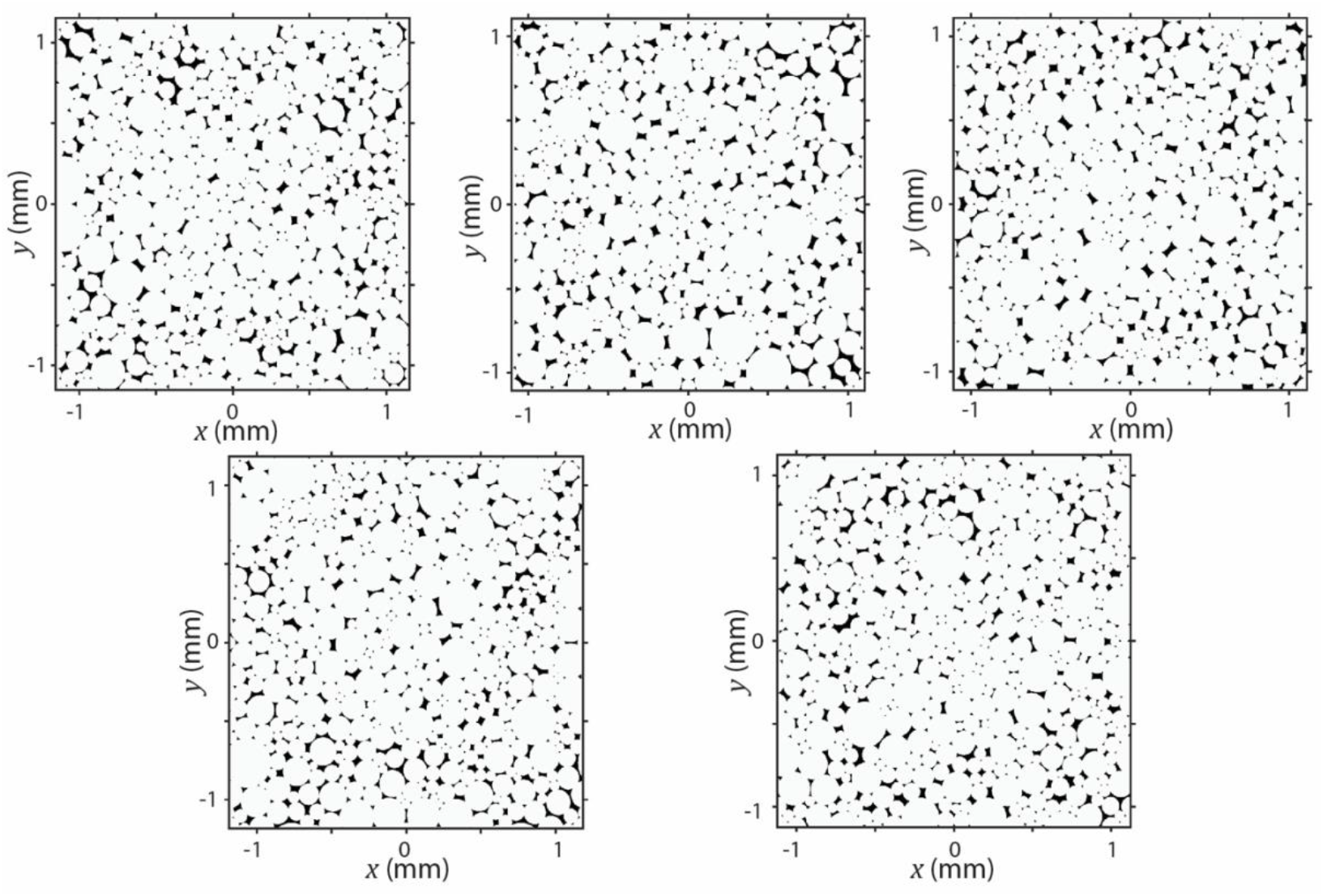
Different 2D REV realizations used to simulate sponge porosity.

**Supplementary figure 7.**
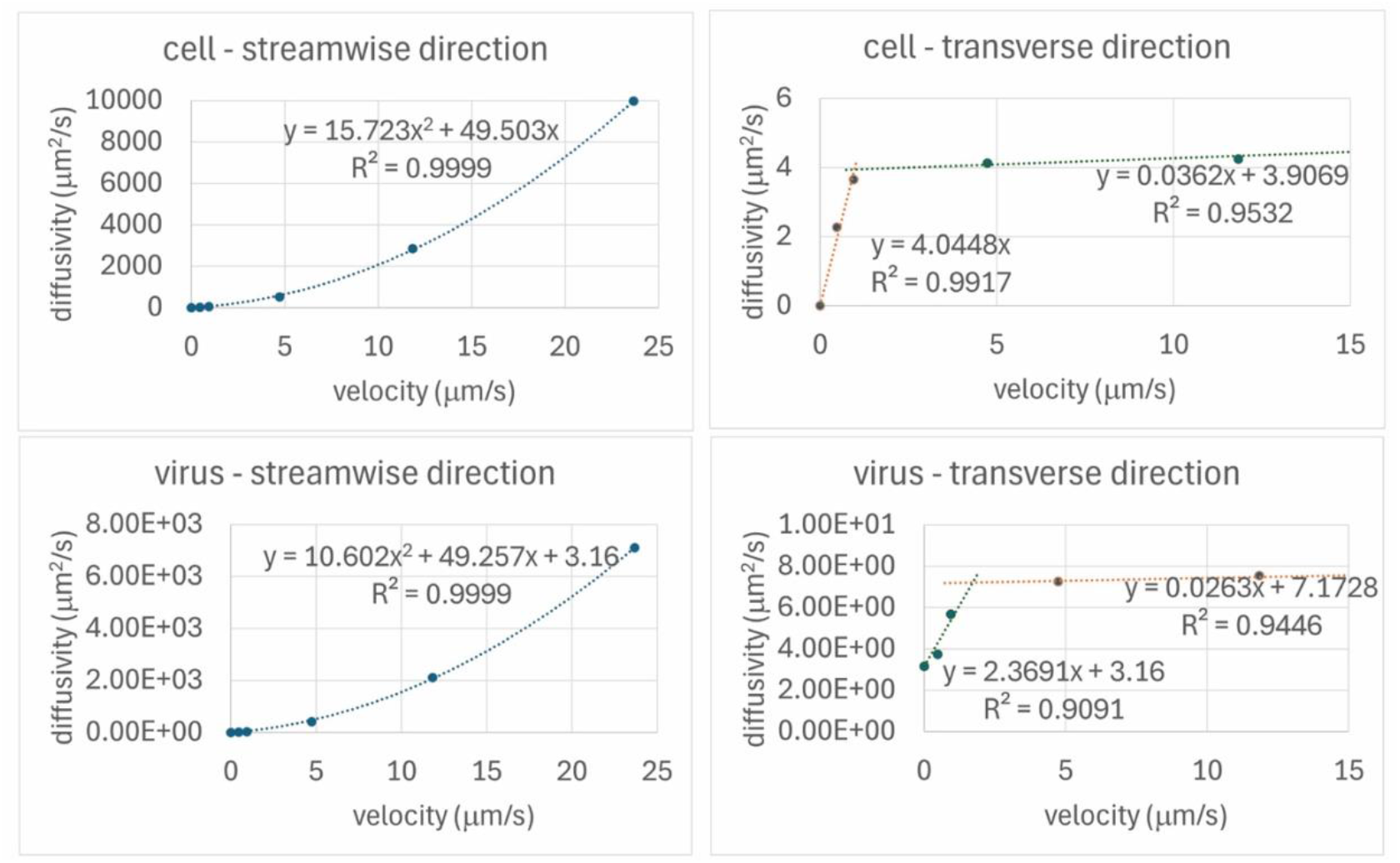
Polynomial fits for effective diffusivity of cells and viruses at different flow velocities.

**Supplementary figure 8.**
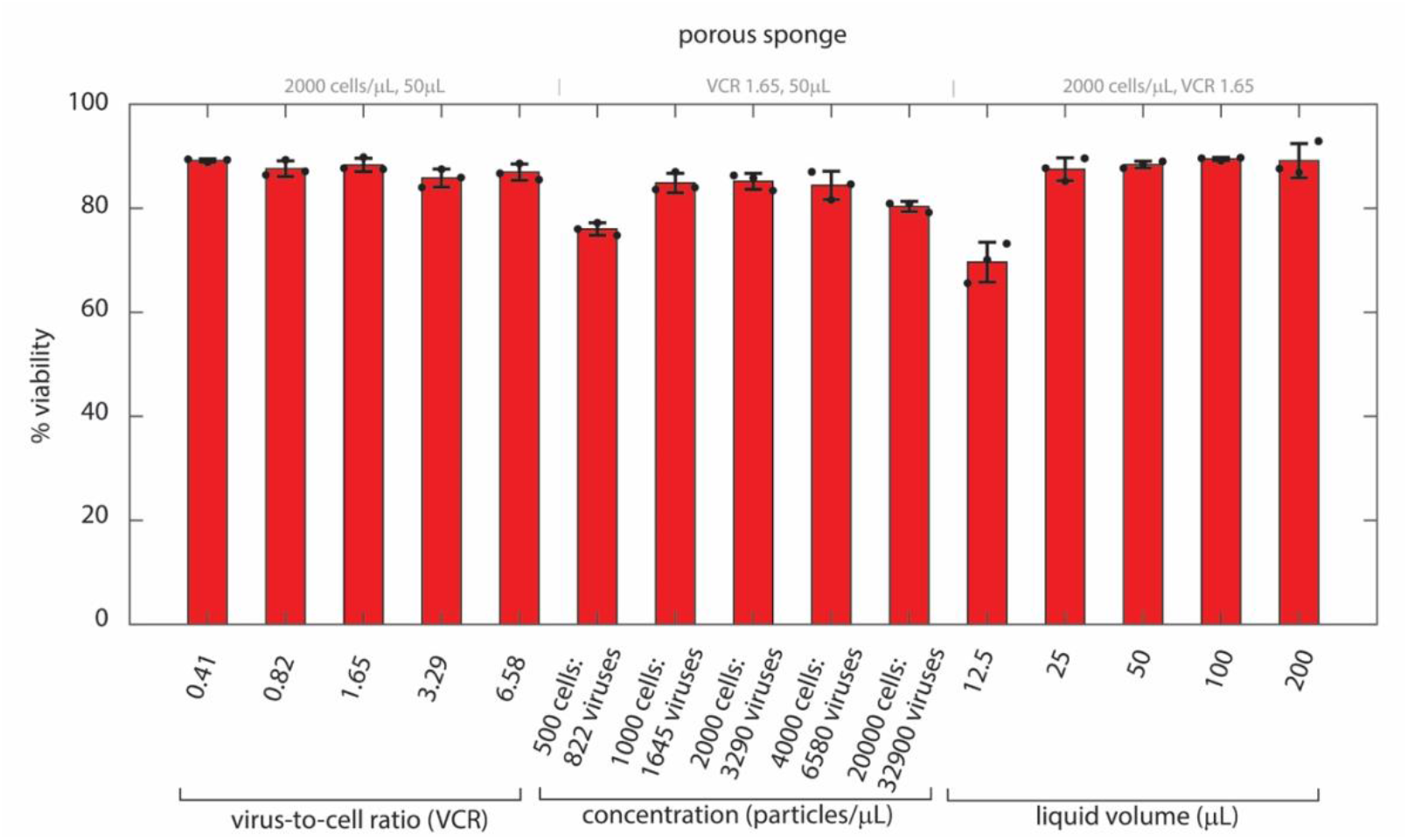
Experimentally measured cell viability in the porous sponge for different VCRs, particle concentrations, and liquid volumes. Cell viability was analyzed after transduction protocols and 72h cell culture.

**Supplementary figure 9.**
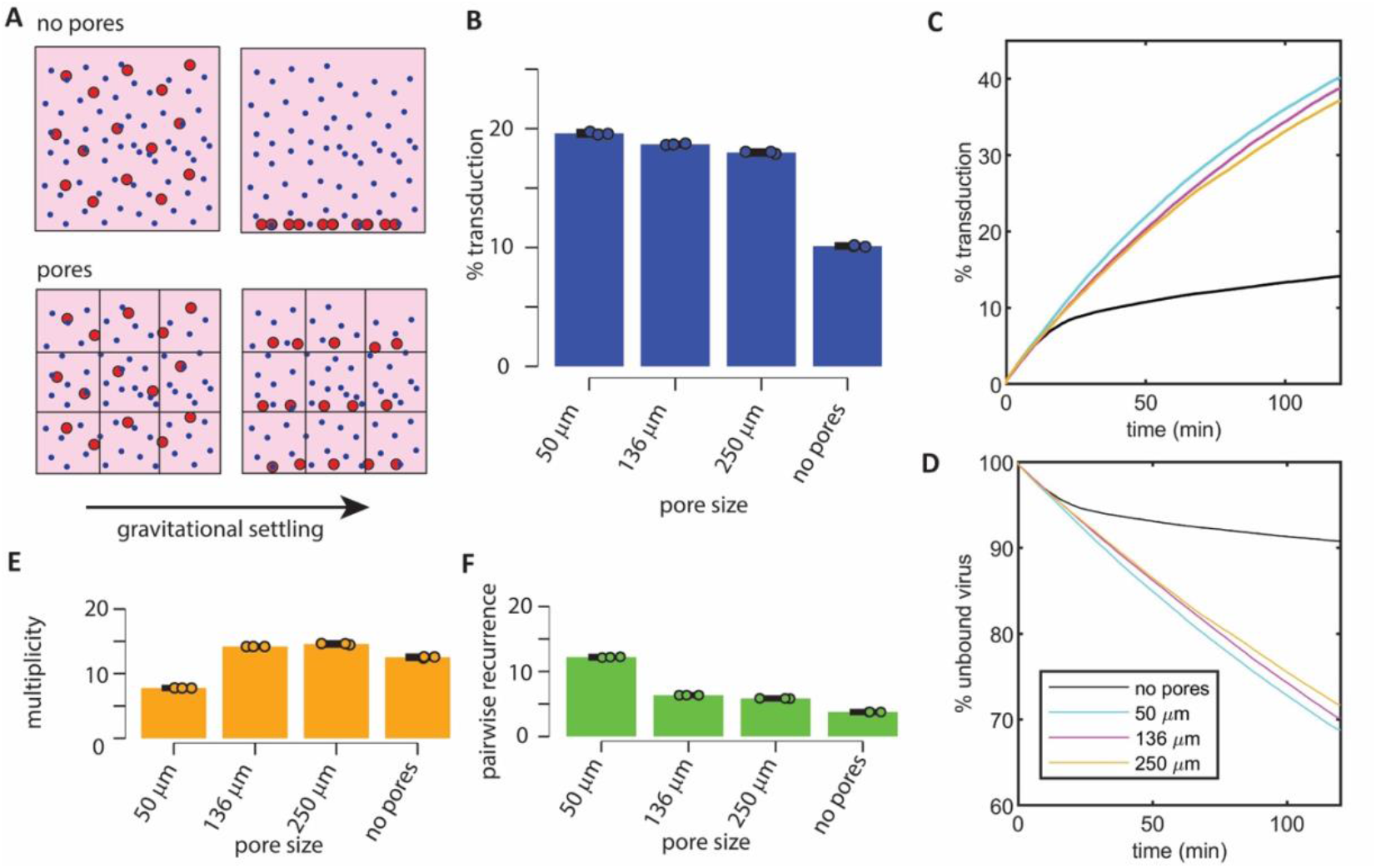
(a) Simulation case investigating the role of particle colocalization revealed that (b) a maximum of 2x transduction improvement can be expected due to this mechanism alone, which could only account for °25% of sponge transduction enhancement. One-way ANOVA of the simulated transduction efficiency confirmed significant differences in transduction efficiency between the no pores and with pore colocalization groups as well as significant differences between different pore sizes (p < 0.001). (c) Pore colocalization led to consistent transduction enhancement over time when compared to the no pore group, (d) which led to improved virus utilization. (e-f) pore colocalization led to increased cell-virus collision frequency with increasing pairwise recurrence of collisions at small pore sizes. Cell-virus concentration = 20000 cells/µL, VCR = 1.65.

**Supplementary figure 10.**
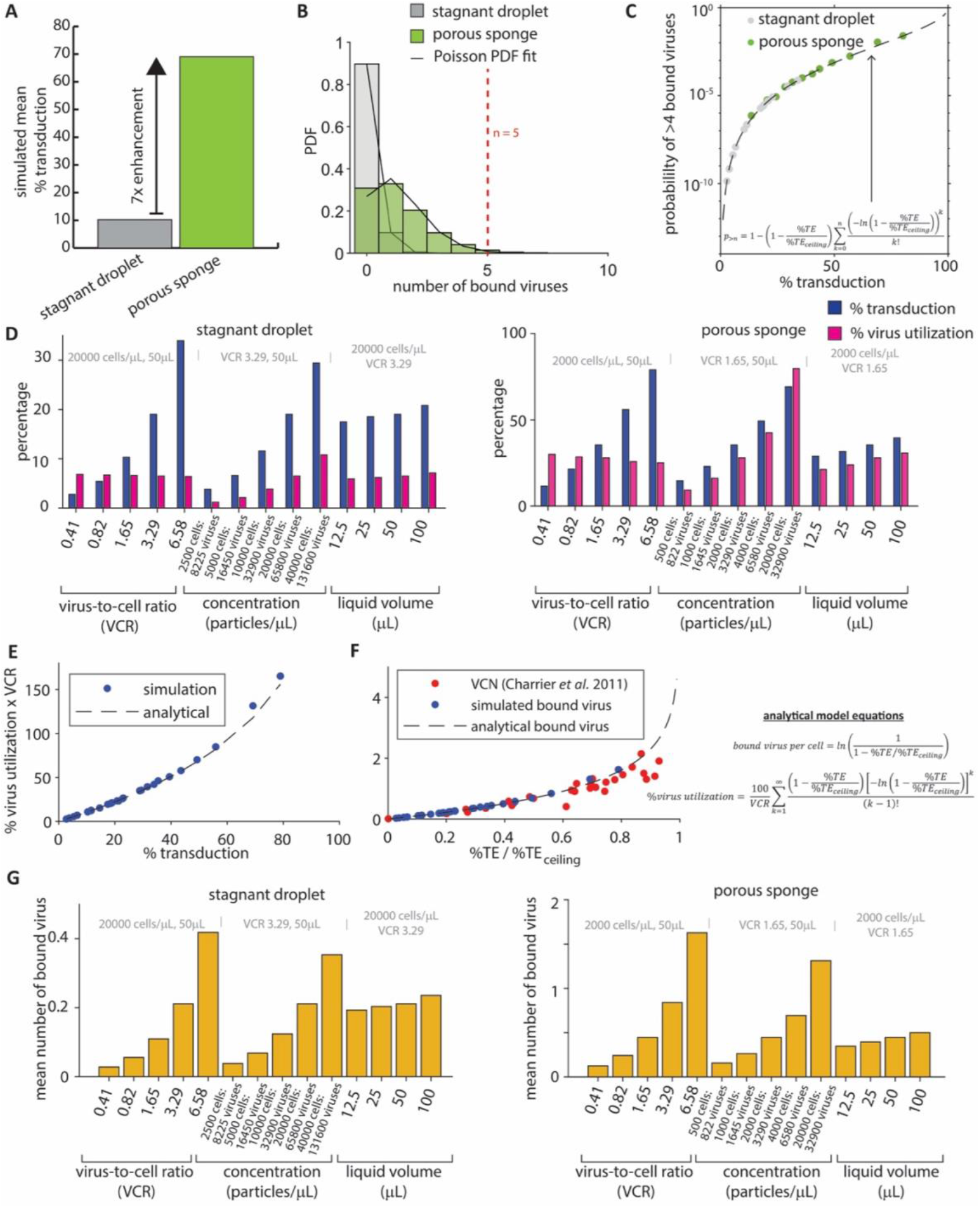
(a) Fold change of simulated transduction efficiency in porous sponges when compared to a stagnant liquid droplet under the same conditions shows multifold transduction improvement in porous sponges (20000 cells/µL, VCR 1.65, 50µL). (b) Probability density distributions of the number of bound viruses for the modeled cells transduced in stagnant droplets and porous sponges (20000 cells/µL, VCR 1.65, 50µL). (c) Predicted probability of cells binding with 5 copies of the virus or more in stagnant droplets and porous sponges. The circle markers show simulated data, and the dashed line shows the analytical curve fit. (d) Viral particles are more effectively utilized in porous sponges when compared to stagnant droplets since high virus utilization at a given VCR is determined by transduction efficiency (e). (f) The mean number of bound viruses per cell is also linked to transduction efficiency leading to higher number of bound viruses per cell in sponges when compared to stagnant droplets (g). The term (*%TE / %TE*_*ceiling*_) is interpreted as the fraction of the maximum transduction capacity of the system where *%TE*_*ceiling*_ is the biological ceiling of transduction efficiency saturation, estimated by VCR titration as 97.400% in our Jurkat-GFP-retrovirus system and 65.649% in the experimental reference using a CD34^+^-GFP-lentivirus system^59^.

**Supplementary figure 11.**
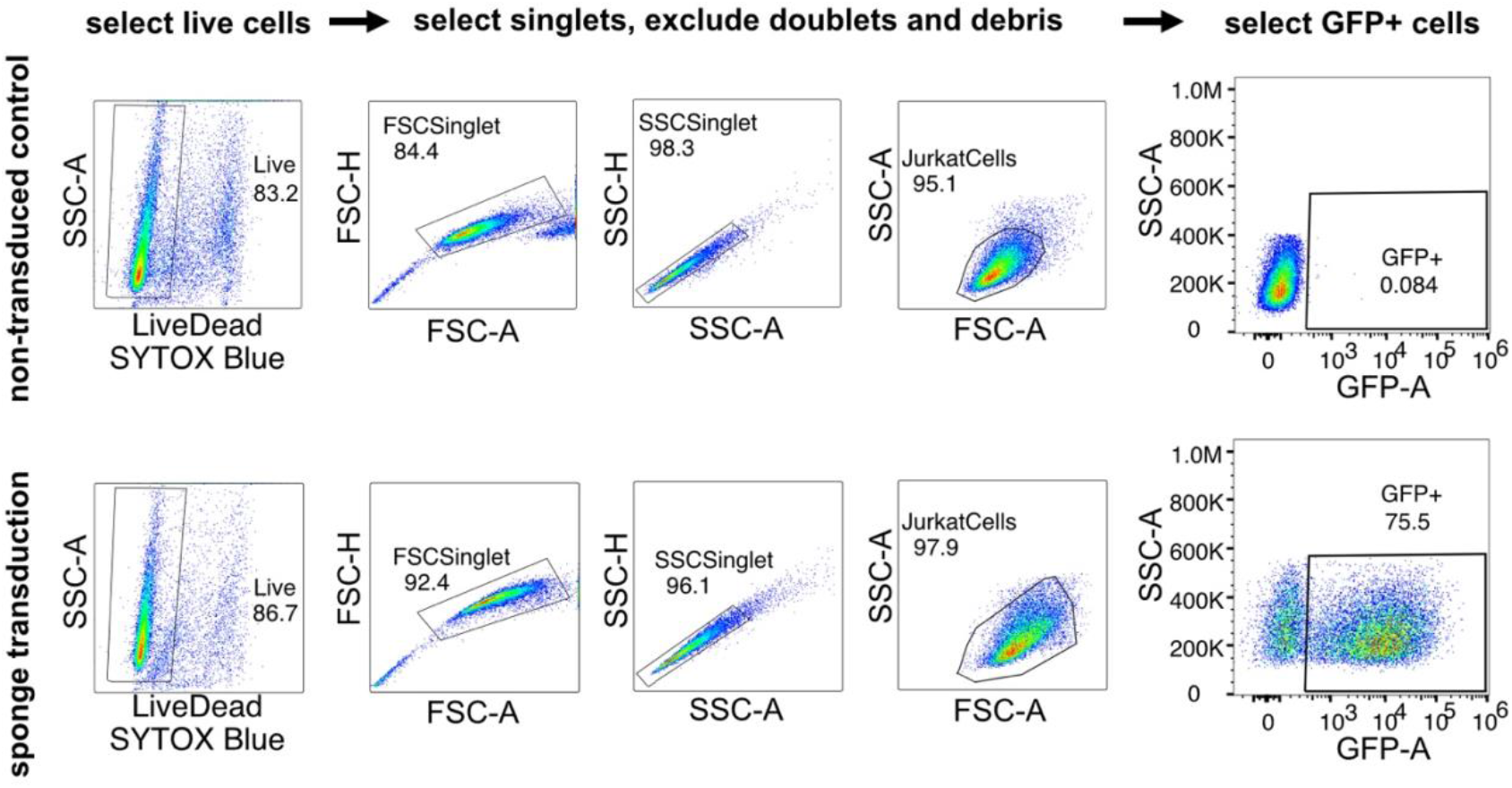
Flow cytometry gating strategy used to analyze GFP+ cells shown for the cases of non-transduced control and transduction in the porous sponge.

## Supplementary tables

**Supplementary table 1:** Summary of cases for transduction in a stagnant liquid droplet. The shaded row represents the case with identical conditions between stagnant droplet and porous sponge.

| case number | liquid volume (μL) | cell concentration (cells/μL) | VCR | droplet height (mm) | mean %TE (simulated) | mean %TE (experiment) |
| --- | --- | --- | --- | --- | --- | --- |
| 1 | 50 | 20,000 | 0.41 | 2.3 | 2.768 | 3.783 |
| 2 | 50 | 20,000 | 0.82 | 2.3 | 5.352 | 6.756 |
| 3 | 50 | 20,000 | 1.65 | 2.3 | 10.251 | 10.933 |
| 4 | 50 | 20,000 | 3.29 | 2.3 | 19.120 | 17.766 |
| 5 | 50 | 20,000 | 6.58 | 2.3 | 34.119 | 28.966 |
| 6 | 50 | 2,500 | 3.29 | 2.3 | 3.863 | 7.153 |
| 7 | 50 | 5,000 | 3.29 | 2.3 | 6.745 | 10.400 |
| 8 | 50 | 10,000 | 3.29 | 2.3 | 11.763 | 13.900 |
| 9 | 50 | 20,000 | 3.29 | 2.3 | 19.120 | 17.700 |
| 10 | 50 | 40,000 | 3.29 | 2.3 | 29.520 | 24.000 |
| 11 | 12.5 | 20,000 | 3.29 | 1.5 | 17.648 | 14.700 |
| 12 | 25 | 20,000 | 3.29 | 1.8 | 18.427 | 16.533 |
| 13 | 50 | 20,000 | 3.29 | 2.3 | 19.120 | 19.233 |
| 14 | 100 | 20,000 | 3.29 | 2.9 | 20.741 | 18.566 |

**Supplementary table 2:** Simulated effective diffusivities of cells and viruses at different flow velocities.

| velocity<br>($\mu\text{m/s}$) | cell diffusivity ($\mu\text{m}^2/\text{s}$) | | virus diffusivity ($\mu\text{m}^2/\text{s}$) | |
| --- | --- | --- | --- | --- |
|  | transverse<br>directions (x, z) | streamwise<br>direction (y) | transverse<br>directions (x, z) | streamwise<br>direction (y) |
| 0 | 0 | 0 | 3.16 | 3.16 |
| 0.473 | 2.264 | 12.903 | 3.729 | 11.098 |
| 0.947 | 3.655 | 36.985 | 5.679 | 32.233 |
| 4.73 | 4.135 | 517.368 | 7.254 | 418.528 |
| 11.8 | 4.247 | 2837.254 | 7.554 | 2112.022 |
| 23.7 | 4.796 | 9971.204 | 7.769 | 7100.727 |

**Supplementary table 3:** Summary of cases for transduction in a porous sponge. The shaded row represents the case with identical conditions between stagnant droplet and porous sponge.

| case number | experimental liquid volume ($\mu\text{L}$) | cell concentration (cells/$\mu\text{L}$) | VCR | droplet height (mm) | Mean %TE (simulated) | Mean %TE (experiment) |
| --- | --- | --- | --- | --- | --- | --- |
| 1 | 50 | 2000 | 0.41 | 2.3 | 11.517 | 18.400 |
| 2 | 50 | 2000 | 0.82 | 2.3 | 20.617 | 30.000 |
| 3 | 50 | 2000 | 1.65 | 2.3 | 35.602 | 45.600 |
| 4 | 50 | 2000 | 3.29 | 2.3 | 56.107 | 68.433 |
| 5 | 50 | 2000 | 6.58 | 2.3 | 79.600 | 79.566 |
| 6 | 50 | 500 | 1.65 | 2.3 | 13.908 | 22.800 |
| 7 | 50 | 1000 | 1.65 | 2.3 | 23.224 | 35.933 |
| 8 | 50 | 2000 | 1.65 | 2.3 | 35.602 | 48.366 |
| 9 | 50 | 4000 | 1.65 | 2.3 | 49.419 | 56.666 |
| 10 | 50 | 20000 | 1.65 | 2.3 | 69.044 | 78.633 |
| 11 | 12.5 | 2000 | 1.65 | 1.5 | 29.012 | 40.866 |
| 12 | 25 | 2000 | 1.65 | 1.8 | 31.412 | 45.833 |
| 13 | 50 | 2000 | 1.65 | 2.3 | 35.602 | 46.866 |
| 14 | 100 | 2000 | 1.65 | 2.9 | 39.503 | 52.400 |

## Supplementary Appendix A Functional viral titer estimation using probabilistic modeling

### Reduced-order transduction model for titer estimation

In this paper, we are considering transduction using porous sponges and stagnant droplets across various cell-virus concentrations and liquid volumes. A very small proportion of the viral vectors in the stock solution contain the machinery to bind to the cell and then subsequently integrate its genetic material into the cell genome. This creates the concept of physical and functional viral titer representing the total number of viral particles and the number of infectious viral particles, respectively. Estimating the functional viral titer accurately is crucial for the quantitative modeling approach in this paper. Therefore, we want to estimate a functional viral titer using the experimentally measured transduction efficiencies that is valid across the entire range of cell and virus concentrations, as well as across two different transduction methods. Crucially, we do not want to make any assumptions about the physical mechanism driving transduction in these different methods so that we do not create biases in the subsequent analyses we perform by using the estimated functional titer value.

Let the number of cells and number of viruses in 1µL of the liquid medium be denoted by *n*_*cell*_ and *n*_*virus*_, respectively. In an ideal scenario where there are infinitely many cell-virus collisions, the probability that a virus transduces a specific cell (*P*_*occupied*_) can be calculated by equation A1.

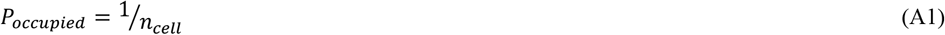

In practical application, transduction is dependent on several rate-limiting upstream processes such as convective-diffusive particle dynamics that lead to cell-virus collisions, the probability that the cell-virus collision leads to viral-receptor binding on the cell-membrane, and probability of successful integration. Therefore, we assume that the *P*_*occupied*_ is also dependent of the probability that these upstream processes occur (*P*_*upstream*_).

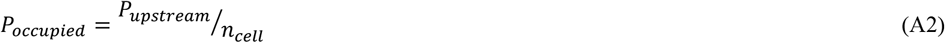

The probability that no viruses transduce the cell can be calculated as 1-*P*_*occupied*_. We can then calculate the probability that every virus avoids transducing a cell as:

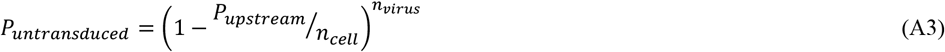

Hence, transduction efficiency (%TE) can be calculated as:

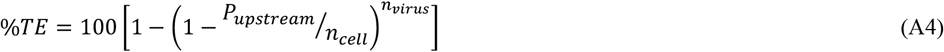

We assumed that *P*_*upstream*_ is a saturating exponential function (equation A5) that depends on the cell concentration – a parameter that controls the separation distance between the cells and determines process efficiency. When *n*_*cell*_ → 0, *P*_*upstream*_ → 0, correctly predicting that the infinitely large separation distance between the cells would lead to zero transduction. When *n*_*cell*_ → ∞, *P*_*upstream*_ → 1, correctly predicting that at high cell concentrations, the distance between the particles is virtually zero and the upstream processes are not the limiting factor in transduction. We also experimentally measured that the transduction ceiling at high VCR is over 98%. This further justified the use of the saturating exponential function without additional scaling factors to consider transduction ceiling.

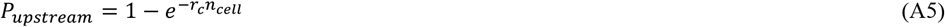

where *r*_*c*_ is the capture rate coefficient.

### Titer estimation using Markov Chain Monte Carlo

We used slice sampling Markov Chain Monte Carlo (MCMC) to estimate the posterior probability distributions of the base virus concentration (*c*_*vb*_, where *c*_*vb*_ *= n*_*virus*_/dilution), the capture rate coefficient (*r*_*c*_), and the standard deviation of the residual between the probabilistic model and experiment (*stdev*). The prior probability distributions are assumed to be equal to 1 in the intervals *c*_*vb*_ ∈ (1,∞), *r*_*c*_ ∈ (0,1), and *stdev* ∈ (0,10). Both *c*_*vb*_ and *r*_*c*_ are explored in exponential steps for robust parameter estimation and uncertainty quantification. First, we applied the MCMC algorithm to estimate the posterior probability distribution of *c*_*vb*_, *r*_*c*_, and *stdev* for the experimentally measured transduction efficiencies in sponge transduction (**supp. figure 12**). We estimated the functional viral titer as the median value of the probability distribution, 3.47 × 10^7^ TUs/mL (95% CI [3.21 × 10^7^, 4.43 × 10^7^]), which yielded a root mean square error between the model and experiment of 5.96%.

**Supplementary figure 12.**
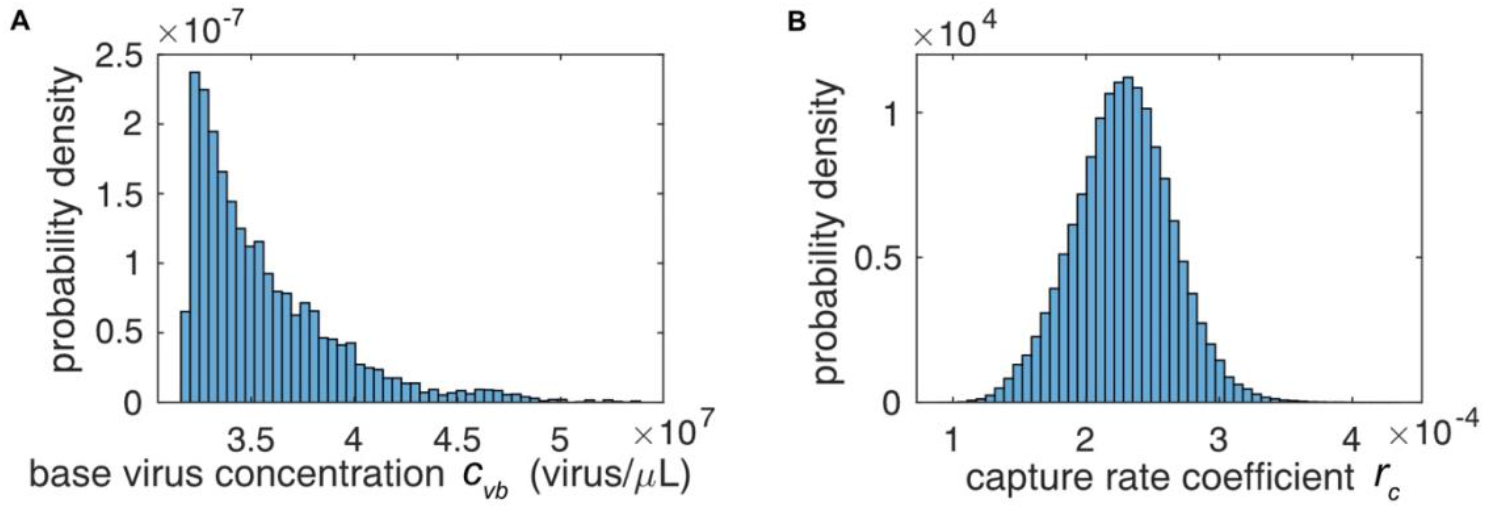
Posterior probability distributions of (a) base virus concentration (*c*_*vb*_) and (b) capture rate coefficient (*r*_*c*_) estimated by MCMC for sponge transduction.

Next, we repeated the analysis for the experimentally measured transduction efficiency in stagnant droplets. The parameter estimation of functional viral titer using the low transduction efficiencies observed in stagnant droplets creates an ill-posed problem. In stagnant liquid droplets, the capture rate coefficient (*r*_*c*_) is significantly smaller when compared to highly efficient methods like porous sponges. Using Taylor series expansion, we can show:

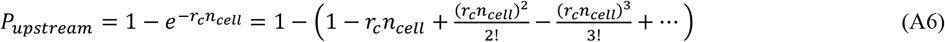

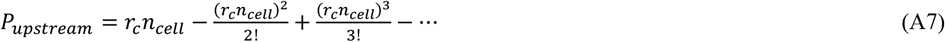

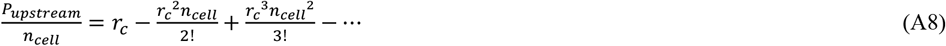

For small *r*_*c*_, such as in the stagnant droplet case, high order terms are omitted and the transduction efficiency can be approximated as:

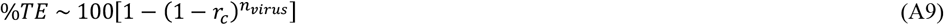

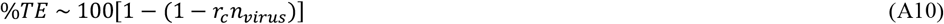

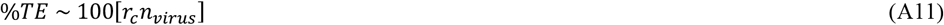

Therefore, at low transduction efficiencies, the problem is ill-posed leading to a non-unique solution and the MCMC algorithm will produce multimodal posterior distributions. To overcome this limitation, we leverage prior knowledge of *c*_*vb*_ from the posterior probability distribution calculated for transduction in porous sponges and assume a prior probability distribution of 1 for *c*_*vb*_ ∈ [3.21 × 10^7^, 4.43 × 10^7^], which is the 95% credible interval of *c*_*vb*_ in sponges. The prior probability distributions are assumed to be equal to 1 in the intervals *r*_*c*_ ∈ (0,1), and *stdev* ∈ (0,10). The MCMC algorithm then estimated the posterior probability distribution using experimentally measured transduction efficiencies in stagnant droplets. We estimated the functional viral titer as the median value of the probability distribution, 3.33 × 10^7^ TUs/mL (95% CI [3.24 × 10^7^, 3.5 × 10^7^]), which yielded a root mean square error between the model and experiment of 3.82%. We then took the average of the functional titers obtained for porous sponges and stagnant droplets (3.40 × 10^7^ TUs/mL) to set up the simulation model.

## Supplementary Appendix B Governing equations of the fluid flow and particle models

### Equilibrium liquid droplet shape model

To calculate the dimensions of the equilibrium liquid droplet shape, we minimize the free-energy of the liquid-air-solid system (*E*) by using equations B1-B4 assuming that the droplet is an ellipsoid cap^30^.

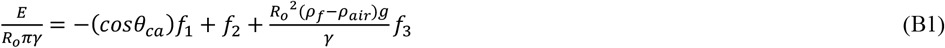

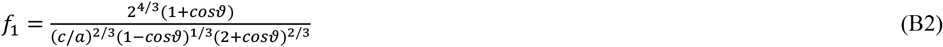

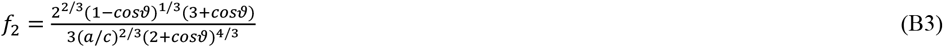

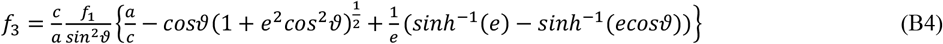

where γ is the surface tension (0.072 N/m), *θ*_*ca*_ is the contact angle (68°), *R*_*o*_ is the radius of an imaginary spherical droplet of the same volume, ρ_*f*_ is fluid density, *ρ*_*air*_ is air density, and *g* is gravitational acceleration. We used a brute force grid search algorithm to find the values of *c*/*a* and ϑthat minimize *E*. Using these values of *c*/*a* and ϑ, we calculate the droplet height *h* and radius *R* using equations B5-B8.

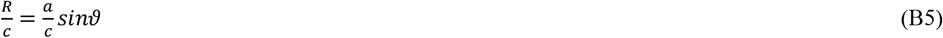

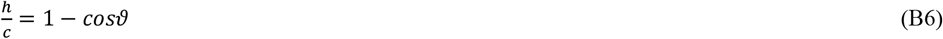

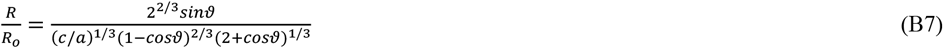

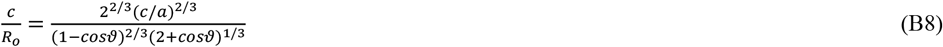

### Liquid absorption model

The governing equation of the 1-D liquid absorption model is written in equation B9-B10, derived from the Young-Laplace and Darcy law equations.

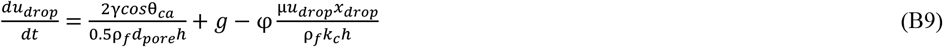

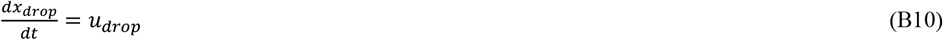

where *d*_*pore*_ is mean pore diameter, *k*_*c*_ is permeability, φ is porosity, and µ is fluid viscosity. The fluid viscosity is assumed to be unchanged by the cell suspension. At the base cell concentration used for the sponge experiments (2000 cells/µL), the effective fluid viscosity is only 1.003 times the molecular viscosity according to Lundgren's formula. At a cell concentration of 20000 cells/µL, the effective fluid viscosity is 1.03 times the molecular viscosity. The governing equations are solved using a semi-implicit time advancement algorithm to predict the position (*x*_*drop*_) and velocity (*u*_*drop*_) of the liquid droplet in the sponge. Markov Chain Monte Carlo (MCMC) using the slice sampling algorithm is used to estimate the posterior probability distribution of *k*_*c*_. The MCMC algorithm treats both *k*_*c*_ and the standard deviation of the residual between simulation and experiment as parameters to be estimated since the standard deviation of the residuals is not known *a priori*. To define the prior probability distribution, *k*_*c*_ and standard deviation values in the range 0 to infinity are assumed valid. The posterior probability distribution estimated by MCMC is then analyzed to calculate the mean value of the *k*_*c*_, which is used in the absorption model, and the 95% credible interval is calculated to quantify model uncertainty.

### Computational Fluid Dynamics model

The Navier-Stokes equations written in equations B11-B12 are solved using the Finite Volume Method (FVM).

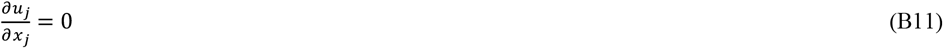

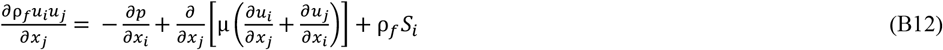

where *u* is fluid velocity, *p* is periodic pressure, and *x* is Eulerian position. The momentum source term *S*_*i*_ is used to specify a linear pressure drop to sustain flow in the periodic domain. The value of *S*_*i*_ required to sustain a flow with a given Reynolds number is not known *a priori*. For incompressible flow, the specification of the mass flow rate is sufficient to maintain a constant superficial velocity through the porous medium. Therefore, *S*_*i*_ is determined iteratively during the pressure correction step from the difference between the desired and the current mass flow rate. The spatial derivatives are approximated using a second-order upwind scheme for the convective terms and a second-order central scheme for the viscous terms. Pressure interpolation on the element face is performed with second order accurate methods. The governing equations are solved in a segregated manner using a SIMPLE algorithm.

The numerical grid used to simulate the flow is generated using the triangulation algorithm in ANSYS Meshing. A grid convergence study was undertaken to conclude that refining the grid beyond a resolution of 10µm resulted in less than 1% improvement in the predicted drag force and mean square streamwise velocity fluctuation. An REV size convergence study was undertaken to conclude that changes in the predicted drag force and mean square streamwise velocity fluctuation are not statistically significant when the number of pores was changed from 25, 50, and 100 pores. The shear stress distribution was analyzed at *u*_*m*_ = 25µm/s to conclude that the maximum shear stress and pressure inside the porous sponge were 0.046 N/m^2^ and 0.063 N/m^2^, whereas the reported values of stress required for mechanoporation are greater than 1000 N/m^2^.

### Discrete Particle model

The equations of motion of the cell and virus particles^67^ are defined in equation B13, which is expanded into equation B14.

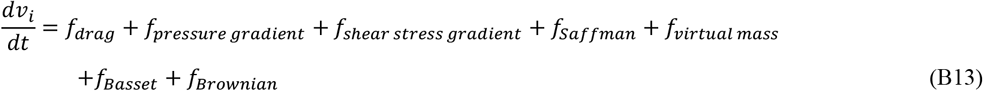

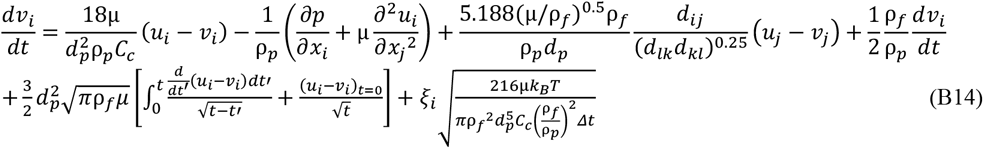

This full model was implemented to calculate a budget of forces acting on the cell and virus particles (*n* = 1,000) flowing through the 2D REV at *u*_*m*_ = 25µm/s for 1,000 seconds. For cells, *f*_*drag*_ and *f*_*pressure gradient*_ were the dominant forces, whereas the remaining forces were at least 3 orders of magnitude smaller, whereas *f*_*drag*_ and *f*_*Brownian*_ were dominant forces for viruses. Based on this insight, we simplified the discrete particle model to equation B15 by modeling only *f*_*drag*_, *f*_*Brownian*_, and *f*_*pressure gradient*_.

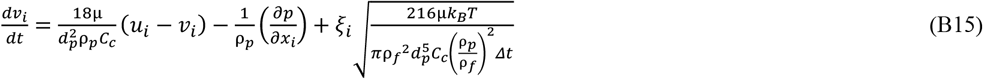

where *v* is particle velocity, *d*_*p*_ is particle diameter, ρ_*p*_ is particle density, *t* is time, *C*_*c*_ is the Cunningham correction factor, ξ is a Gaussian white noise generator, *T* is temperature, Δ*t* is time step, and *k*_*B*_ is the Boltzmann constant. We assumed that the system was isothermal since all the components are held at 37°C. While evaporative cooling can theoretically induce Marangoni convection even in isothermal environments, our experiments were conducted in a humidified incubator (>95% RH), which minimizes evaporation-driven surface tension gradients. Furthermore, any weak convective mixing arising from such effects would be insignificant in comparison to gravitational settling and Brownian motion.

The particle equations of motion are solved using Euler implicit discretization for stable time advancement. Time step sensitivity of the solver was verified for time step sizes ranging between 0.0001s and 0.1s to confirm that all of these time step values predict the same particle trajectory. At the macroscale level, gravity is modeled throughout the simulation volume considering buoyancy and an additional effective diffusivity force (*f*_*effective diffusivity*_) is included in the model where porosity is less than 1.

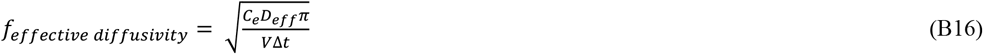

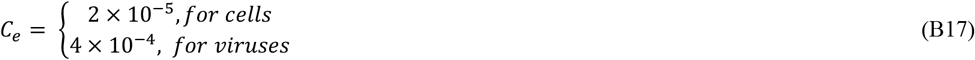

where *V* is the particle volume. The model constant *C*_*e*_ is calibrated to simulate the correct particle diffusivity with a model fit of R^2^>0.99. Time step size is changed according to the particle diffusivity (Δ*t* = 0.1*cell diameter/2*D*_*eff*_). This ensures that cell-virus collisions are not missed in between time steps. Physically, this also implies that when the virus collides with a cell, it will diffuse over a constant value of the cell surface area in between finite time steps in the model. This is crucial to compare how particle dynamics across different cases translate towards the same objective of finding a receptor on the cell membrane to bind and leading to successful transduction^14^.

### Transduction Model

At the end of the simulation, the collision history between cells and viruses are recorded, saving the time stamp, cell and virus IDs, position of the collision event. Each cell-virus collision event is translated into a transduction event with two possible outcomes - transduced or not transduced, determined by the probability of transduction (*p*_*TE*_). Transduction events are determined by simulating a Bernoulli trial assuming probability of success of *p*_*TE*_ using a uniformly distributed random number generator (function rand() in MATLAB). Following this approach, the transducing collisions are analyzed to calculate a probability density distribution of bound viruses over all of the cells. The simulated transduction efficiency is then calculated as the number of cells with greater than zero bound viruses divided by the total number of cells.

The model is first calibrated using cell-virus collision histories for transduction in the stagnant liquid droplet for the variation of the particle concentration. Sponge simulation data is not included in this calibration to prevent model bias towards high transduction efficiencies observed in the sponge, allowing fair evaluation of the mechanisms of collisions in sponge transduction. To define the prior probability distribution, *p*_*TE*_ values in the range 0 to 1 are assumed valid and standard deviation values in the range 0 to infinity are assumed valid. The MCMC algorithm is used to estimate the posterior probability distribution of *p*_*TE*_ (**supp. figure 13**). The mean value of the *p*_*TE*_, 0.0023 (95% CI [0.0010,0.0043]), is then used in the transduction model to predict transduction efficiency.

**Supplementary figure 13.**
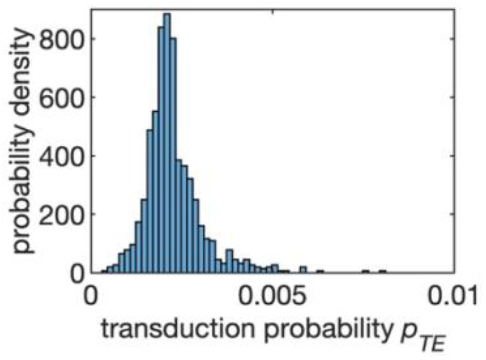
Posterior probability distributions of transduction probability (*p*_*TE*_) estimated by MCMC for transduction in a stagnant liquid droplet for different cell concentrations.

## Notes

### Competing Interest Statement

The authors declare the following financial interests/personal relationships which may be considered as potential competing interests: Y.B, has pending patents related to CAR T cell production and therapy. Y.B. is a scientific founder of Persistence Therapeutics which seeks to translate technology related to this work. Y.B. serves on the Board of Persistence and owns equity in Persistence Therapeutics. Y.B. has licensed technology for reagents to transduce cells to Takara Bio USA. Y.B. receives an industry-sponsored research grant related to CAR T cell therapeutic technology unrelated to this work.

